# Gut feeling: host and habitat as drivers of the microbiome in blackbuck (*Antilope cervicapra*)

**DOI:** 10.1101/2024.02.20.581148

**Authors:** Ananya Jana, Shamik Roy, Sumanta Bagchi, Kavita Isvaran, K Praveen Karanth

## Abstract

The gut microbiome can be shaped by both intrinsic host factors and extrinsic environmental factors. However, the relative importance of intrinsic and extrinsic factors in gut microbial composition has rarely been investigated, particularly for a single host across its natural range. Here, we characterise the gut microbiome of an endemic, endangered antelope, the blackbuck or *Antilope cervicapra*. We evaluated the influence of seven predictor variables, which were classified into intrinsic and extrinsic factors, on the gut microbiome. We determined which of these seven variables explains greater variation in the microbiome within (α-diversity) and between (β-diversity) the blackbuck host. We analysed the microbiome of 60 blackbuck hosts from ten different populations across India. We recorded 11800 unique OTUs across 30 known phyla and 2.9 million reads. We find an average of 2056 OTUs per individual, with firmicutes and bacteroidetes being the most dominant phyla. We find that nucleotide diversity (intrinsic), blackbuck population density (intrinsic), distance to human settlement (extrinsic), and anthropogenic land-use (extrinsic) explain within-host variation. In contrast, precipitation (extrinsic), nucleotide diversity, distance to human settlement, and anthropogenic land-use explain between-host variation. Overall, we also show that the genetic diversity of the host is more important than their environment for both within- and between-host variation in the microbiome, in blackbuck. Therefore, conservation efforts should be directed to not only preserve natural habitats but also increase the genetic pool of the blackbuck populations, which will positively impact their survival through diverse gut microbiomes.

## Introduction

The gut microbiome is highly dynamic, and much progress has been made in understanding how intrinsic (e.g., host diversity) and extrinsic (e.g., environment and geography) factors regulate the gut microbiome. For instance, there exists a phylogenetic inertia where host phylogeny across different species and/or genetic relatedness among individuals in a species determines gut microbiome structure, especially in mammals (Amato et al., 2018; Ley et al., 2008; Muegge et al., 2011; Sanders et al., 2015; Song et al., 2020; Youngblut et al., 2019). In other words, genetically related species/individuals tend to harbour similar gut microbiomes. These patterns of microbiome clustering with host phylogeny and genetic diversity can be via ‘vertical transmission’ that includes germline colonization, infant contact with maternal microbiota during birth and physical contact between parents and their progeny, or via ‘horizontal transmission’ that includes interactions of host with their conspecifics and environment (Mallott & Amato, 2021). Separately, extrinsic factors such as climate and anthropogenic pressure can also shape the gut microbiome. Indeed, under a laboratory-based experimental setup, a 2-3 °C rise in environmental temperature altered the gut microbiota with negative consequences for the host (Bestion et al., 2017). It was also found that varying degrees of anthropogenic activities can affect gut microbiota (Knutie et al., 2019; Lavrinienko et al., 2021). Thus, intrinsic and extrinsic factors can independently impact the gut microbiome. However, it remains unclear in wild populations whether the effect of intrinsic host factors on gut microbiota outweighs that of extrinsic environmental factors or vice versa (Kartzinel et al., 2019; Knutie et al., 2019). It also remains unknown whether the relative importance of extrinsic and intrinsic factors changes with gut microbiome within individual hosts (i.e., α-diversity of the microbiome) or between individuals of the host population (i.e., β-diversity of the microbiome). A comparative study of the gut microbiome of a single species across its geographic range can allow us to resolve yet unanswered questions. Identifying and elucidating essential ecological determinants that shape the gut microbiome in a host can help understand the links between gut microbiota and its host, unravel climate-induced shifts in the microbiota that, in turn, can influence host fitness, and formulate conservation management plans to protect host populations.

Resolving the relative importance of extrinsic and intrinsic factors for gut microbiome is an ongoing challenge that has been impeded by multiple reasons. First, several studies were performed on captive animals rather than wild populations, which has hindered our comprehensive understanding of the potential determinants of the gut microbiome (Alberdi et al., 2021; Bornbusch et al., 2022; Nelson et al., 2013; Schmidt et al., 2019; Weinstein et al., 2021). It is now well known that the gut microbiome of captive animals substantially differs from their wild counterparts, and therefore they might not represent the true natural processes (McKenzie et al., 2017). Second, studies on mammalian gut microbiomes have focussed on phylogenetically diverse species sets where intra-species variation in host diversity has largely been ignored (Amato et al., 2018; Knowles et al., 2019; Youngblut et al., 2019). This can introduce bias in causal inferences because a few individuals from each species might not capture true intra-species variation. Third, following the last reason, most studies have been conducted on a single host population rather than different host populations across wide-ranging geographies (Davies et al., 2022; Murillo, Schneider, Fichtel, et al., 2022; Stoffel et al., 2020). This can obscure the genomic and environmental underpinnings of the observed differences in the gut microbiome.

Here, we attempt to overcome these challenges by analysing gut bacterial communities at different scales (α- and β-diversity) in an endemic, endangered antelope, the blackbuck or *Antilope cervicapra,* across its current range. Blackbuck are sexually dimorphic, even-toed, medium-sized antelope (23–45 kg; Mungall, 1978; Ranjitsinh, 1989) that belong to the Bovidae family, phylogenetically nested within the gazelle clade (Jana & Karanth, 2023, 2019). The monotypic genus is a very recently diverged lineage, only ∼2 million years ago (Jana & Karanth, 2019) and provides us with a model system to examine the effects of anthropogenic changes on their gut microbiota while accounting for phylogenetic constraints. The charismatic animal is a flagship of the grassland ecosystem, substantial parts of which have been razed following urbanization and expansion of agriculture, and therefore, blackbuck is often a direct target of human-wildlife conflict. Since grasslands in India remain heavily fragmented and are under immense pressure for land-use change, threatening the long-term survival of blackbuck, it is timely and imperative to ascertain the determinants of the blackbuck’s gut microbiome. This will help us understand how blackbuck respond and/or adapt to the drastically changing environment and draft effective conservation management strategies.

The intrinsic host factors considered were blackbuck population size and genetic diversity (see Table 1). The latter was assessed both in terms of nuclear (microsatellite) and mitochondrial markers in order to obtain a comprehensive view of the genetic structure in this model system, as the two classes of biomarkers can be influenced by different aspects of their evolutionary history (Jana & Karanth, 2023). While both these markers evolve rapidly in animal systems and can provide information on recent population genetic changes, mitochondrial DNA is primarily inherited maternally (Moore, 1995). Hence, differences in animal behaviour between the sexes with respect to the mating system and philopatry, among others, can lead to incongruency between the patterns observed in genetic diversity from different classes of genetic markers. Extrinsic environmental factors were classified into human influence and climate. Human influence was quantified in two ways: 1. distance between the blackbuck population and closest human settlements, and 2. percentage of the land cover influenced by humans (agriculture, settlements and plantations) that the blackbuck population occupies or, in other words, anthropogenic land use (see Table 1). Climatic variables were temperature and precipitation (see Table 1).

**Table 1:** Predictor variables and their importance for gut microbiome.

| Variables | Importance |
| --- | --- |
| **Precipitation** | Variation in rainfall and quality of water sources can lead to differences in resource availability and foraging and feeding behaviour, thus correlating with differences in gut microbiome (Murillo, Schneider, Fichtel, et al., 2022). |
| **Temperature** | Climate warming is a threat to all life forms and the change in ambient temperature can result in loss of susceptible microbial species and/or increase in the more resistant pathogenic microbes (Bestion et al., 2017). |
| **Population density** | Host density could be correlated with physical contacts or social interaction within the species and hence affect microbial transmission (Li et al., 2016). |
| **Nucleotide diversity** | Unique traits (e.g., viviparous birth, lactation, co evolution of the placenta and the immune system) of the host animal can lead to patterns of microbiome composition being affected by host phylogeny (Mallott & Amato, 2021). Nucleotide diversity in our study considers the maternally derived mitochondrial genetic information. |
| **Heterozygosity** | Unique traits (e.g., viviparous birth, lactation, co-evolution of the placenta and the immune system) of the host animal can lead to patterns of microbiome composition being affected by host phylogeny (Mallott & Amato, 2021). Heterozygosity will account for genetic composition of the host that is derived from both maternal and paternal sources. |
| **Distance to Human settlement** | Sites with varying human presence can change the environment and diet (like introducing novel food items) of the host animal, which in turn can alter the gut microbiota (Knutie et al., 2019). |
| **Anthropogenic land-use** | Anthropogenic influences, like change in land-use can alter both food sources and environmental microbiome, which directly affect the composition of the gut microbiome (San Juan et al., 2020). |

Here, we ask two interdependent questions: 1. Whether and how each of the seven predictor variables (between extrinsic and intrinsic factors) influence the gut microbiome of blackbuck within an individual host (α-diversity) and between different individuals (β-diversity)? 2. What explains greater variation in the α- and β-diversity of blackbuck gut microbiome, intrinsic host or extrinsic environmental factors? Based on a priori literature available on gut microbiome from other hosts, we expected blackbuck gut microbiome to vary with population size (H. Li et al., 2016), nuclear diversity (Mallott & Amato, 2021), heterozygosity (Mallott & Amato, 2021; Stoffel et al., 2020), distance to human settlements (Knutie et al., 2019), anthropogenic land use (San Juan et al., 2020), temperature (Bestion et al., 2017; Kikuchi et al., 2016), and precipitation (Murillo, Schneider, Fichtel, et al., 2022). However, we did not have any expectations of the direction and magnitude of the effects of each variable on the microbiome. Given that the environment-host-microbe interactions are complex, and the environment and host can independently determine microbiome structure, the overall relative effect of the host (intrinsic) and habitat (extrinsic) related factors in this particular model system remains to be unravelled.

## Materials and Methods

### Study system

Our focal system is the Indian blackbuck or *Antilope cervicapra*, a monotypic genus endemic to the Indian subcontinent (Fig. S1). Apart from the chinkara, the endemics viz., nilgai, the four-horned antelope and the blackbuck are the only antelopes found in India. However, only *A. cervicapra* forms a part of the tribe of ‘true antelopes’ found in the world (Jana & Karanth, 2019). Adult males are characterized by their dark dorsal colouration, white underbelly, and striking spiral horns, which are permanent. The females are light brown in colour and do not have horns. They are group-living grazers and mainly subsist on a grass-dominated diet. Although a flagship of the semi-arid grassland ecosystem, they seem well adapted to open woodlands and even scrublands. Categorized as a Schedule I (endangered) species under the Wildlife Protection Act of India, 1972, blackbuck have come under intense pressure in the past due to hunting and poaching and, more recently, due to habitat loss and fragmentation. Currently, relegated to multiple small patches across India (many of them isolated), blackbuck populations show much variation in population density, proximity to human habitation, and diet, among others.

### Sampling

The fecal samples of blackbuck were collected from 10 locations across their geographic range. The sampling was conducted at Bhetanoi, Haliya, Tal Chhappar, Velavadar, Nannaj, Ranebennur, Kolar, Timbuktu, Rollapadu, and Point Calimere. All collections were done non-invasively, without animal handling, and no animals were disturbed in the process. Blackbuck were observed from a distance and tracked on foot or followed by vehicles. As diurnal animals, blackbuck generally forage during the early morning and afternoon, and hence, the sampling was mostly conducted during this period. The GPS coordinates were recorded at each collection location. The population densities were estimated from field observations and personal communications from surveyors and forest department reports (Jana & Karanth, 2023). As we had access to more accurate density records rather than the actual number of individuals in any area, the densities worked as proxies for population sizes for the purpose of our study. Blackbuck fecal pellets were easily differentiated from those of other sympatric bovids due to their characteristic fecal shape (Bhaskar et al., 2021), and removed any possibility of any potential mix-up. Fresh fecal samples from adult blackbuck individuals were taken, where whole pellets were collected and stored in 100% ethanol. The samples were stored at room temperature when in the field site. Once transferred to the laboratory, the DNA extraction procedure was carried out.

### DNA Extraction

DNA was extracted from the whole pellets using the QIAmp Fast Stool Extraction kits (Qiagen, Germany) following the standard protocols after select modifications. After scraping the outer layer of the pellets into a fine powder, the samples were submerged in an extraction buffer overnight before we proceeded to the next steps. Post extraction, the samples were quality checked using a nanodrop and Qubit fluorimeter and stored at -20 °C.

### Genetic analyses

#### Nucleotide diversity

The D loop region of the mitochondrial genome was amplified to estimate nuclear diversity (Jana & Karanth, 2023, 2019). Primers were designed de novo using the sequence for the D loop region obtained from an *A. cervicapra* genome deposited in GenBank (Accession Code: AO003422.1). Following preliminary trials, four primer pairs namely HV1D, INT, DLF3 and DLF4 were shortlisted, where each pair ∼300 bp of the D loop region and together covered the entire ∼ 800 bp of the D loop region (Jana & Karanth, 2023). The successfully amplified PCR products for every primer pair and each sample were sent to our registered vendors, Medauxin and Barcode Biosciences (Bangalore, India), for Sanger sequencing. Every sample was sequenced in both forward and reverse directions.

The final sequence obtained was aligned with the GenBank data to remove the risk of potential ‘numts’ (nuclear copies of mitochondrial DNA) and ensure the accuracy of the mitochondrial DNA region of blackbucks. To correct any errors in the identification of the nucleotide bases, the obtained sequences were edited and cleaned using Chromas v2.6.5 (technelysium.com.au/ chromas.html). Cleaned sequences were then aligned using the Muscle Algorithm with default parameters in Mega v7 (Tamura et al., 2011). After the final clean-up and alignment of the complete sequences, the nucleotide diversity values for each of the sampling locations were calculated using DnaSP v6.11. 01 (Rozas et al., 2017). Briefly, nucleotide diversity was calculated as follows: first, for a pair of complete D loop sequences that represent two separate individuals, the frequency of nucleotide differences between the paired sequences was determined. Second, the calculated frequency was divided by the total length (in bp) of the paired sequences to estimate nucleotide differences per base position for the pair. Third, previous steps were repeated for all the possible pairs in a given sampling location. Fourth, the nucleotide diversity was calculated as the average of the nucleotide differences per base position across all the pairs of sequences in the population. *See additional methods for more details*.

#### Heterozygosity

Heterozygosity of the blackbuck population was estimated following genotyping (Jana & Karanth, 2023, 2019). Genotyping is the method to identify the alleles of each microsatellite. Following preliminary analyses, only seven primer pairs: bm302, bm415, maf70, hdz496, inra040, tgla122 and sps115 (Jana & Karanth, 2023) were selected for this study. These primers showed positive amplification in more than half of the tested samples in the preliminary trials. The final PCR products were sent to Barcode Biosciences (Bangalore, India) with LIZ500 as the size standard to generate an electropherogram for each locus and individual.

Each locus for every sample was amplified separately at least three times to keep allele identification errors (scoring errors) at a minimum. For all replicates, the resulting electropherograms were compared against each other for consistent peaks. For each sample, microsatellite alleles at each of the 7 loci (corresponding to the 7 primer pairs) were identified manually upon visual inspection of the electropherogram. Here, individuals having two differently sized alleles are heterozygous for that microsatellite locus, while those with the same allele are homozygous. Further quality checks were done to reduce genotyping errors and to increase confidence in our results. Quality checks were done using MICROCHECKER v2.2.3 (Van Oosterhout et al., 2004) to detect the presence of null alleles and allelic drop-out, and using GenAlEx v6.503 package (Peakall & Smouse, 2006, 2012) in Microsoft Excel 2016 to calculate Pid/PI and Pid_sibs_/PI_sibs_ values. From these quality checks, one can conclude that the set of seven microsatellite loci were correctly genotyped. Following all the quality checks, population-wise unbiased observed heterozygosity was calculated. Heterozygosity (Ho) is the proportion of heterozygous alleles in each population. Population-wise heterozygosity was calculated in two steps: first, the proportion of heterozygote loci was calculated for each individual. For example, if an individual is heterozygous for 3 out of 7 loci, then the proportion would be 3/7, which is 0.429. Second, the average of the proportion values for all individuals in the population was considered population-wise heterozygosity. *See additional methods for more details*.

### Gut microbiome

Around 25 ng of the DNA was used to amplify the hypervariable V3-V4 region of 16S rRNA of the bacterial genome using modified 341F (5’-CCTACGGGNGGCWGCAG-3’) and 785R (5’- GACTACHVGGGTATCTAATCC-3’) primers. KAPA HiFi HotStart Ready Mix was used for polymerase chain reactions, involving an initial denaturation step at 95 °C for 5 minutes, followed by 25 cycles of 95 °C for 30 seconds, 55 °C for 45 seconds and 72 °C for 30 seconds and a final extension at 72 °C for 7 minutes. The amplicons were purified using Ampure beads to remove unused primers. Additionally, 8 cycles of PCR were performed using Illumina barcoded adapters to prepare the sequencing libraries. The library was quantified using Qubit DNA HS quantitation assay (Thermo Scientific), which specifically quantifies double-stranded DNA assay. The library was then taken for further sequencing. The adapter sequences used were the P7 adapter for read 1 (AGATCGGAAGAGCACACGTCTGAACTCCAGTCA) and the P5 adapter for read 2 (AGATCGGAAGAGCGTCGTGTAGGGAAAGAGTGT). The sequencing was outsourced to Clevergene Biocorp Pvt. Limited, based in Bangalore, India. The sequence data was generated using Illumina MiSeq, and the data quality was checked using FastQC and MultiQC software. The data was checked for base call quality distribution and sequence adapter contamination, and all the samples cleared the QC threshold (Q20>95%).

Mothur Ver 1.43.0 (Schloss et al., 2009) was used on a Linux platform to run the bioinformatic pipeline to obtain the final processed data from the MiSeq output. The raw reads were summarized into contigs sequentially trimmed to a uniform length. Data from all the samples were rarefied (rarefaction curves were drawn for each sample) to verify that saturation had occurred and that the number of reads obtained per sample was sufficient to proceed with our analyses. The unique sequences were filtered, the chimeric sequences were removed, and the data was categorized into clusters. The SILVA database (Quast et al., 2013) was used as a reference to align the contig clusters and build the final operational taxonomic unit (OTU) tables. The OTU information was further used to construct a phylogeny which was then used for downstream analyses with R programming language.

### Predictor variables

The seven chosen factors (Table S1) were broadly classified into a) abiotic factors i.e., temperature and precipitation; b) host (blackbuck) related factors i.e., population density, nucleotide diversity and observed heterozygosity; and c) human-mediated factors i.e., distance to closest human settlement and percentage of human-influenced land cover.

The average temperature and precipitation data at each sampling location was obtained from the Climate Research Unit database (CRU v4.03) with a resolution of 0.5^0^ (∼55 km at the equator) for the sampling month from 2014 to 2019. The nucleotide diversity and heterozygosity values were obtained from the genetic analyses as mentioned earlier. The details of landcover types were categorised following the classification provided by BHUVAN using Linear Imaging Self-scanning Sensor (LISS-3) satellite imagery in a area of ∼75 square km with sampling location at the centre. Agricultural lands, settlements, orchards and mixed plantations were considered human-influenced land types. Water bodies were excluded from the calculation of the area covered by these land use types. Human influence on blackbuck was calculated as the distance to the nearest human settlement from the sampling location of the blackbuck, and anthropogenic land-use as a fraction of total land cover by agricultural lands, settlements, orchards and mixed plantations in 75 km^2^ area around each blackbuck population.

### Data analyses

Blackbuck gut microbiome diversity was summarised as α-diversity and β-diversity. These were calculated for each individual fecal sample, representing an individual blackbuck’s gut microbiome. α-diversity includes species richness and Shannon diversity, and β-diversity includes dispersion (distance to centroid) and the first 3 axes of Principal Coordinates Analysis (PCoA) based on the Bray-Curtis dissimilarity matrix. Species richness takes only the number of species present into account, while the Shannon index also incorporates the evenness of the community. Dispersion is a multivariate measure of heterogeneity in species composition, estimated as the distance of each sample relative to their overall group centroid in a multivariate space. Species richness, Shannon diversity, and dispersion were estimated using the ‘*vegan*’ library in R 4.1.1 (Oksanen et al., 2019). PCoA was performed using the ‘*ape*’ library in R 4.1.1 (Paradis et al., 2004). Species diversity (α- and β-diversity) were response variables, and abiotic factors (precipitation, temperature), host factors (population density, nucleotide diversity, heterozygosity), and human factors (distance to human settlement, anthropogenic land-use) were predictor variables.

#### Univariate analyses

Five models were fitted that can potentially describe the relationship between blackbuck gut microbiome diversity and each of seven predictor variables: Linear, Exponential, Power, Logarithm (natural log), and Quadratic (Fig. 2, *See additional methods for more details*). For every predictor variable, model fit was checked for each model following the information-theoretic approach (Burnham & Anderson, 2002). First, the best model fit is selected following Akaike information criteria (AIC; Isvaran & Ponkshe, 2013). The threshold for choosing the best model(s) is considered to ΔAIC<2, where any model that is within 2 AIC (Δ) of the best-performing model (lowest AIC) is considered equally likely and explains considerable variation in species diversity. Second, from the pool of models that were within the threshold of ΔAIC<2, parameter estimates and confidence interval (95% confidence intervals, CI) of the modelled predictor variables were assessed. The models for which the 95% CI were consistently positive or negative, i.e., the lower and higher bounds of the 95% CI do not overlap with zero. Inferences on the relationship between the predictor variable and dependent variable (measures of species diversity) were based on AIC and evaluating the parameter estimates and 95% CI. Further, for predictor variables with significant quadratic relationships, it was tested whether the curve obtains the ‘hump’ (highest value of diversity) or ‘pit’ (lowest value) within the range of observed values for the predictor variable. If neither of the ‘hump’ or ‘pit’ is present, then the relationship might not have a significant quadratic term and is therefore considered linear. These were done using the ‘MOStest’ function of the ‘*vegan*’ package in R 4.1.1.

**Figure 1.**
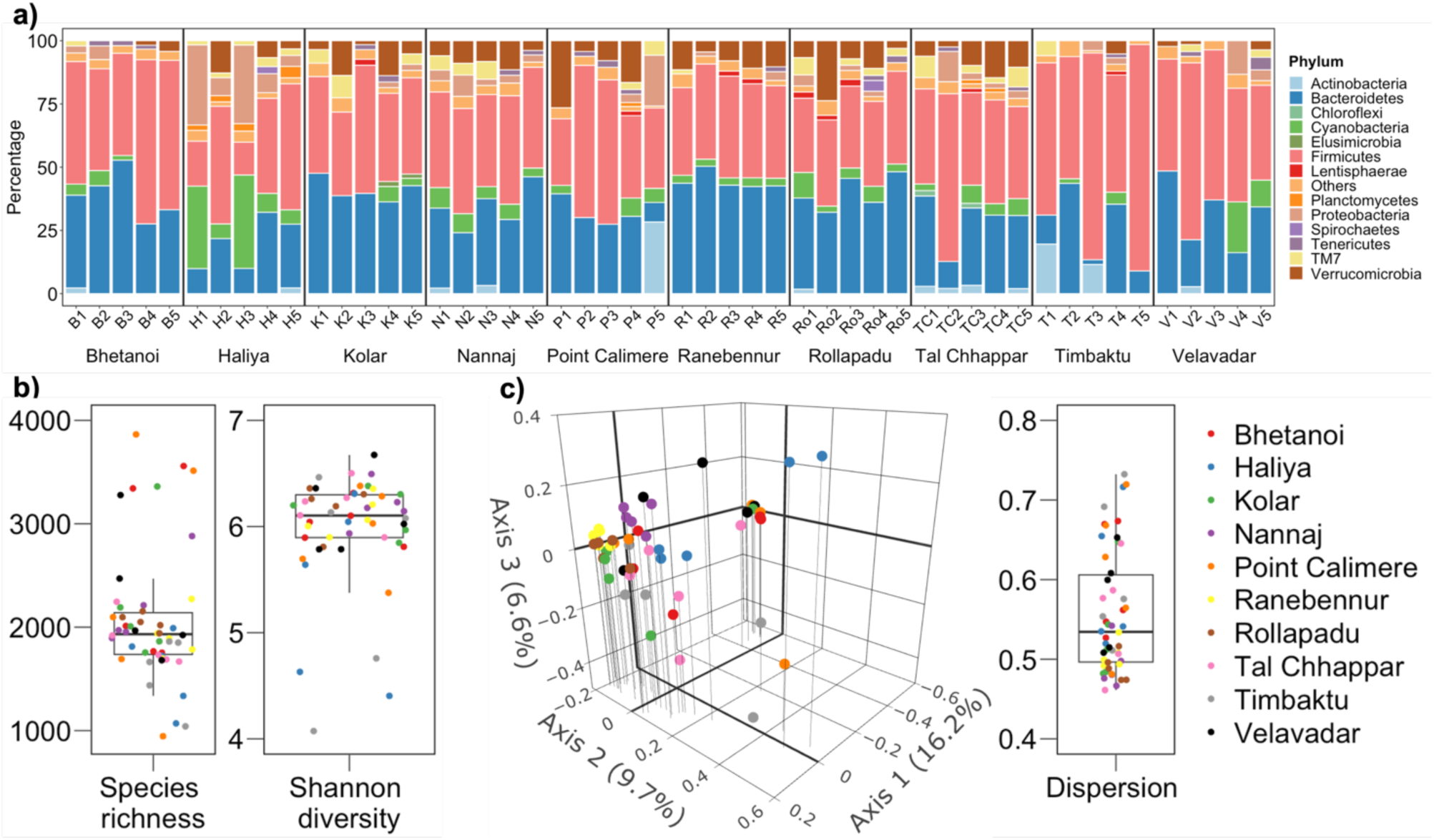
Microbial diversity, a) relative abundance of different microbial phyla across different blackbuck individuals. b) Species diversity at local level (within-individual). c) species diversity at landscape levels summarised as Principal Co-ordinate Analysis (PCoA). PCoA axis 1-3 were taken since together they explain 32.5% of the variation in microbial composition. Dispersion is calculated as distance between individual points and group centroid in a multivariate space (PcoA). In b and d boxplots represent median, IQR and data extremes.

**Figure 2:**
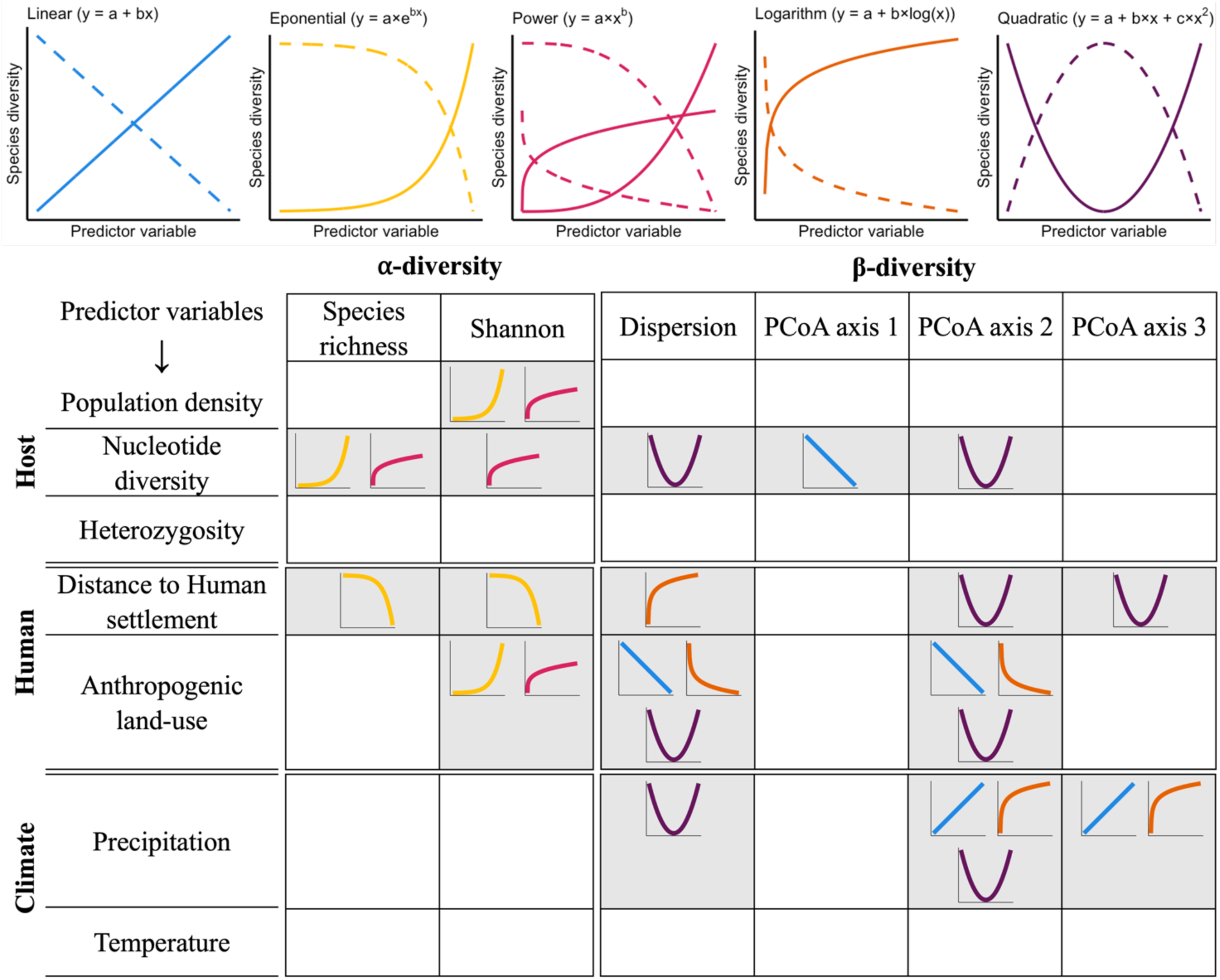
Schematic showing the different models that we fit to microbial species diversity (α-diversity and β-diversity) vs. predictor variables (Climate, Host, Human). Table summarising univariate model fits.

#### Multivariate analyses

Following univariate analysis, multivariate analyses were performed since they helped assess the effect of predictor variables on the species diversity (of the gut microbiome of an individual blackbuck) after evaluating the inter-relationship between all predictor variables. First, a correlation test was performed between all variables to remove highly correlated (r>0.8) variables to reduce multicollinearity in the multivariate models. Temperature and Population density were removed since they were correlated (r>0.8) with Nucleotide diversity (Fig. S3). Therefore, the global models for all species diversity indices (both α- and β-diversity) consist of five predictor variables (fixed-effects): precipitation, nucleotide diversity, heterozygosity, distance to human settlement, and anthropogenic land-use. Second, we built global models following the generalized linear model (GLM) using the ‘*nlme*’ package in R 4.1.1 (Pinheiro & Bates, 2000). Here, the five predictor variables are modelled as fixed-effects, and the sampling location as random-effect to account for the spatial structure in the data (i.e., the samples from multiple individuals from the same population). For every diversity index, multiple global models were created that separately incorporate all the best-performing univariate functions (ΔAIC<2 or ΔAIC=0), such as linear, logarithm, for every predictor variable. For example, two models were examined for species richness, one that modelled nucleotide diversity as an exponential model and another as a power model. But for both models, precipitation was modelled as exponential since that is best-performing univariate function where its ΔAIC=0, even though there is no univariate relationship between species richness and precipitation. Third, similar to univariate analysis, Akaike information criteria was used (AIC; Burnham & Anderson, 2002) to select the best global model(s) for every diversity index. Model(s) with ΔAIC>2 was considered as considerably poorer in performance compared with the best model. Finally, multiple model inference analyses was performed on the best-performing global model(s) for every diversity index. Here, the ‘*MuMIn*’ package in R 4.1.1 generated many sub-models with all combinations of predictor variables in the global model. From the sub-model pool, the best model was chosen with AIC where ΔAIC<2. Model averaged parameter estimates and confidence intervals (averaging across all models within ΔAIC<2) were estimated for inference.

#### Multiple regression on distance matrices (MRM)

Influence of the seven predictor variables on the blackbuck gut microbial community composition was evaluated following a distance-based multiple regression approach using the ‘MRM’ function in the ‘*ecodist*’ package in R 4.1.1 (Goslee & Urban, 2007). MRM, or multiple regression on distance matrices, is an extension of the mantel test and multiple linear regression. Briefly, MRM tests for the association between dissimilarity in species composition and the predictor distance matrices using permutation tests of significance for regression coefficients. Here, microbial community composition was summarised using a pair-wise Bray-Curtis dissimilarity matrix, indicating β-diversity. The genetic distance was accounted for, from both mitochondrial and microsatellite information. The mitochondrial control region sequences were used to calculate the pairwise p-distance, i.e., the number of base differences per site, to generate a distance matrix for nucleotide diversity. From the microsatellite dataset, we obtained the allele size information from each of the loci, which was also used to build a pairwise distance matrix (unbiased Nei’s distance) between individuals. The dissimilarity matrix with Euclidean distance was calculated for precipitation, temperature, population density, distance to human settlement, and anthropogenic land-use. Spatial information (space) on the latitude/longitude of the sampling locations was expressed in Universal Transverse Mercator (UTM) coordinates followed by a dissimilarity matrix with Euclidean distance. A baseline MRM model contains the spatial information from the sampling locations as explanatory variables. This analyzes the role of spatial autocorrelation in determining the patterns in microbial community composition. To this baseline model, we added abiotic, host, and human predictor variables both separately and all together to estimate their influence on microbial community composition.

## Results

### Host and environmental factors

Precipitation (range, 55.0 to 284.7 mm; Fig. S2) and temperature (25.0 to 30.3 °C; Fig. S2) ranges suggested that the sampling locations have a semi-arid climate. There was considerable variation in population density, which ranged from 1 to 150 blackbuck km^-2^. Blackbuck diversity calculated as nucleotide diversity (0.015 to 0.065; Fig. S2) and heterozygosity (0.256 to 0.579; Fig. S2) were also variable. We find that blackbuck can be found in landscapes with almost no influence of humans to wholly dominated by humans since the distance to human settlement ranged between 180 m and 3300 m, and anthropogenic land-use varied from 1.9% to 93.1% (Fig. S2).

### Gut microbiome

We recorded 11800 unique OTUs across 30 known phyla and 2981812 reads. We find that gut microbiota is dominated (Fig. 1, S4) by Firmicutes (mean ± SE, 43.4 ± 2.03%), Bacteroidetes (32.2 ± 1.77%), Cyanobacteria (5.4 ± 0.99%), Proteobacteria (4.3 ± 0.95%) and Verrucomicrobia (6.4 ± 0.84%). Clostridia (Firmicutes, 38.3 ± 2.19%) and Bacteroidia (Bacteroidetes, 32.0 ± 1.82%) were the most abundant classes.

Species richness of OTUs ranged between 945 and 3865 per sample (mean 2056 ± 629 SD; Fig. 1). Shannon diversity ranged between 4.07 and 6.67 per sample (5.98 ± 0.52; Fig. 1). Dispersion (distance to centroid) ranged between 0.46 and 0.73 per sample (0.56 ± 0.08; Fig. 1). A sample here is a fecal sample representing an individual blackbuck.

### Univariate relationship

Overall, the α-diversity of the blackbuck gut microbiome varied with nucleotide diversity (host), blackbuck population density (host), distance to human settlement (human), and anthropogenic land-use (human) (Fig. 3, S5; Table S1, S2). We find that species richness showed a statistically detectable univariate relationship with nucleotide diversity (Exponential model, Power model; ΔAIC<2) and distance to human settlement (Exponential model; ΔAIC<2). Shannon diversity showed a statistically detectable univariate relationship with population density (Exponential model, Power model; ΔAIC<2), nucleotide diversity (Power model; ΔAIC<2), distance to human settlement (Exponential model; ΔAIC<2), and anthropogenic land-use (Exponential model, Power model; *P*=0.05, ΔAIC<2).

Overall, the β-diversity of the blackbuck gut microbiome varied with precipitation (abiotic), nucleotide diversity (host), distance to human settlement (human), and anthropogenic land-use (human) (Fig. 3, S6; Table S1, S2). We find that dispersion showed a statistically detectable univariate relationship with precipitation (Quadratic model; ΔAIC<2), Nucleotide diversity (Quadratic model; ΔAIC<2), distance to human settlement (Logarithm model; ΔAIC<2), and anthropogenic land-use (Linear model, Quadratic model, Logarithm model; ΔAIC<2). Microbial species composition summarised as PCoA axis 1 showed a statistically detectable univariate relationship only with nucleotide diversity (Linear model; ΔAIC<2). Microbial species composition summarised as PCoA axis 2 showed a statistically detectable univariate relationship with precipitation (Linear model, Quadratic model, Logarithm model; ΔAIC<2), Nucleotide diversity (Quadratic model; ΔAIC<2), distance to human settlement (Quadratic model; ΔAIC<2), and anthropogenic land-use (Linear model, Quadratic model, Logarithm model; ΔAIC<2). Microbial species composition summarised as PCoA axis 3 showed a statistically detectable univariate relationship with precipitation (Linear model, Logarithm model; ΔAIC<2) and distance to human settlement (Quadratic model; ΔAIC<2).

### Multivariate relationship

The multivariate model building followed by model simplification showed that blackbuck nucleotide diversity alone best explained the variation in both α-diversity measures, species richness and Shannon diversity (Table 2–3). For all β-diversity measures, we find that blackbuck nucleotide diversity and heterozygosity best explained the variation (Table 2–3).

**Table 2:** Global multivariate mixed effect generalised linear models. Sample location is considered as random-factor for all models. Green indicates the best model (lowest AIC) for any diversity measure.

| Diversity measure | Model No. | Model | AIC | logLik |
| --- | --- | --- | --- | --- |
| Species richness | 1 | $\ln(\text{Species richness}) \sim \ln(\text{Precipitation})$<br>+ *Nucleotide diversity* + $\ln(\text{Heterozygosity})$<br>+ *Distance to Human settlement*<br>+ *Anthropogenic landuse* | 54.74 | -19.37 |
| | 2 | $\ln(\text{Species richness}) \sim \ln(\text{Precipitation})$<br>+ $\ln(\text{Nucleotide diversity})$<br>+ $\ln(\text{Heterozygosity})$<br>+ *Distance to Human settlement*<br>+ *Anthropogenic landuse* | 61.11 | -22.55 |
| Shannon | 1 | $\ln(\text{Shannon}) \sim \text{Precipitation} + \text{Nucleotide diversity} +$<br>$\ln(\text{Heterozygosity}) + \text{Distance to Human settlement} +$<br>$\ln(\text{Anthropogenic landuse})$ | -27.57 | 21.78 |
| | 2 | $\ln(\text{Shannon}) \sim \text{Precipitation} + \ln(\text{Nucleotide diversity}) +$<br>$\ln(\text{Heterozygosity}) + \text{Distance to Human settlement} +$<br>$\ln(\text{Anthropogenic landuse})$ | -35.76 | 25.88 |
| Dispersion | 1 | $\text{Dispersion} \sim \text{Precipitation} + I(\text{Precipitation}^2) +$<br>*Nucleotide diversity* + $I(\text{Nucleotide diversity}^2) +$<br>*Heterozygosity* + $I(\text{Heterozygosity}^2) +$<br>$\ln(\text{Distance to Human settlement}) +$<br>*Anthropogenic landuse* | -60.09 | 41.04 |
| | 2 | $\text{Dispersion} \sim \text{Precipitation} + I(\text{Precipitation}^2) +$<br>*Nucleotide diversity* + $I(\text{Nucleotide diversity}^2) +$<br>*Heterozygosity* + $I(\text{Heterozygosity}^2) +$<br>$\ln(\text{Distance to Human settlement}) +$<br>*Anthropogenic landuse* + $I(\text{Anthropogenic landuse}^2)$ | -39.65 | 31.83 |
| | 3 | $\text{Dispersion} \sim \text{Precipitation} + I(\text{Precipitation}^2) +$<br>*Nucleotide diversity* + $I(\text{Nucleotide diversity}^2) +$<br>*Heterozygosity* + $I(\text{Heterozygosity}^2) +$<br>$\ln(\text{Distance to Human settlement}) +$<br>$\ln(\text{Anthropogenic landuse})$ | -68.36 | 45.18 |
| **PCoA axis 1** | 1 | *PCoA axis 1 ~ Precipitation + Nucleotide diversity + Heterozygosity + I(Heterozygosity<sup>2</sup>) + Distance to Human settlement + ln( Anthropogenic landuse)* | 33.23 | -7.62 |
| **PCoA axis 2** | 1 | *PCoA axis 2 ~ Precipitation + Nucleotide diversity + I(Nucleotide diversity<sup>2</sup>) + ln(Heterozygosity) + Distance to Human settlement + I(Distance to Human settlement<sup>2</sup>) + Anthropogenic landuse* | 28.66 | -4.33 |
|  | 2 | *PCoA axis 2 ~ Precipitation + Nucleotide diversity + I(Nucleotide diversity<sup>2</sup>) + ln(Heterozygosity) + Distance to Human settlement + I(Distance to Human settlement<sup>2</sup>) + Anthropogenic landuse + I(Anthropogenic landuse<sup>2</sup>)* | 46.73 | -12.36 |
|  | 3 | *PCoA axis 2 ~ Precipitation + Nucleotide diversity + I(Nucleotide diversity<sup>2</sup>) + ln(Heterozygosity) + Distance to Human settlement + I(Distance to Human settlement<sup>2</sup>) + ln(Anthropogenic landuse)* | 20.66 | -0.33 |
|  | 4 | *PCoA axis 2 ~ Precipitation + I(Precipitation<sup>2</sup>) + Nucleotide diversity + I(Nucleotide diversity<sup>2</sup>) + ln(Heterozygosity) + Distance to Human settlement + I(Distance to Human settlement<sup>2</sup>) + Anthropogenic landuse* | 49.95 | -13.98 |
|  | 5 | *PCoA axis 2 ~ Precipitation + I(Precipitation<sup>2</sup>) + Nucleotide diversity + I(Nucleotide diversity<sup>2</sup>) + ln(Heterozygosity) + Distance to Human settlement + I(Distance to Human settlement<sup>2</sup>) + Anthropogenic landuse + I(Anthropogenic landuse<sup>2</sup>)* | 67.55 | -21.77 |
|  | 6 | *PCoA axis 2 ~ Precipitation + I(Precipitation<sup>2</sup>) + Nucleotide diversity + I(Nucleotide diversity<sup>2</sup>) + ln(Heterozygosity) + Distance to Human settlement + I(Distance to Human settlement<sup>2</sup>) + ln(Anthropogenic landuse)* | 42.52 | -10.26 |
|  | 7 | *PCoA axis 2 ~ ln(Precipitation) + Nucleotide diversity + I(Nucleotide diversity<sup>2</sup>) + ln(Heterozygosity) + Distance to Human settlement + I(Distance to Human settlement<sup>2</sup>) + Anthropogenic landuse* | 17.58 | 1.21 |
| | 8 | *P*CoA axis 2 ~ $\ln(\text{Precipitation}) + \text{Nucleotide diversity} + I(\text{Nucleotide diversity}^2) + \ln(\text{Heterozygosity}) + \text{Distance to Human settlement} + I(\text{Distance to Human settlement}^2) + \text{Anthropogenic landuse} + I(\text{Anthropogenic landuse}^2)$ | 34.76 | -6.38 |
| | 9 | *P*CoA axis 2 ~ $\ln(\text{Precipitation}) + \text{Nucleotide diversity} + I(\text{Nucleotide diversity}^2) + \ln(\text{Heterozygosity}) + \text{Distance to Human settlement} + I(\text{Distance to Human settlement}^2) + \ln(\text{Anthropogenic landuse})$ | 9.91 | 5.05 |
| **P**CoA axis 3 | 1 | *P*CoA axis 3 ~ $\text{Precipitation} + \text{Nucleotide diversity} + I(\text{Nucleotide diversity}^2) + \ln(\text{Heterozygosity}) + \text{Distance to Human settlement} + \text{Anthropogenic landuse}$ | -14.20 | 16.10 |
| | 2 | *P*CoA axis 3 ~ $\ln(\text{Precipitation}) + \text{Nucleotide diversity} + I(\text{Nucleotide diversity}^2) + \ln(\text{Heterozygosity}) + \text{Distance to Human settlement} + \text{Anthropogenic landuse}$ | -26.50 | 22.25 |

### Multiple regression on distance matrices (MRM)

Distance-based approach with MRM (Table 4) showed that after accounting for all the seven predictor variables, geographical distance (space) between the blackbuck populations and heterozygosity had a statistically detectable effect on the gut microbiota (β-diversity), where space and heterozygosity together explain 6.6% of the variation in the microbiota (F=43.40, *P*<0.001). However, space alone explains 4.8% of the variation (F=62.13, *P*=0.001).

**Table 3:**
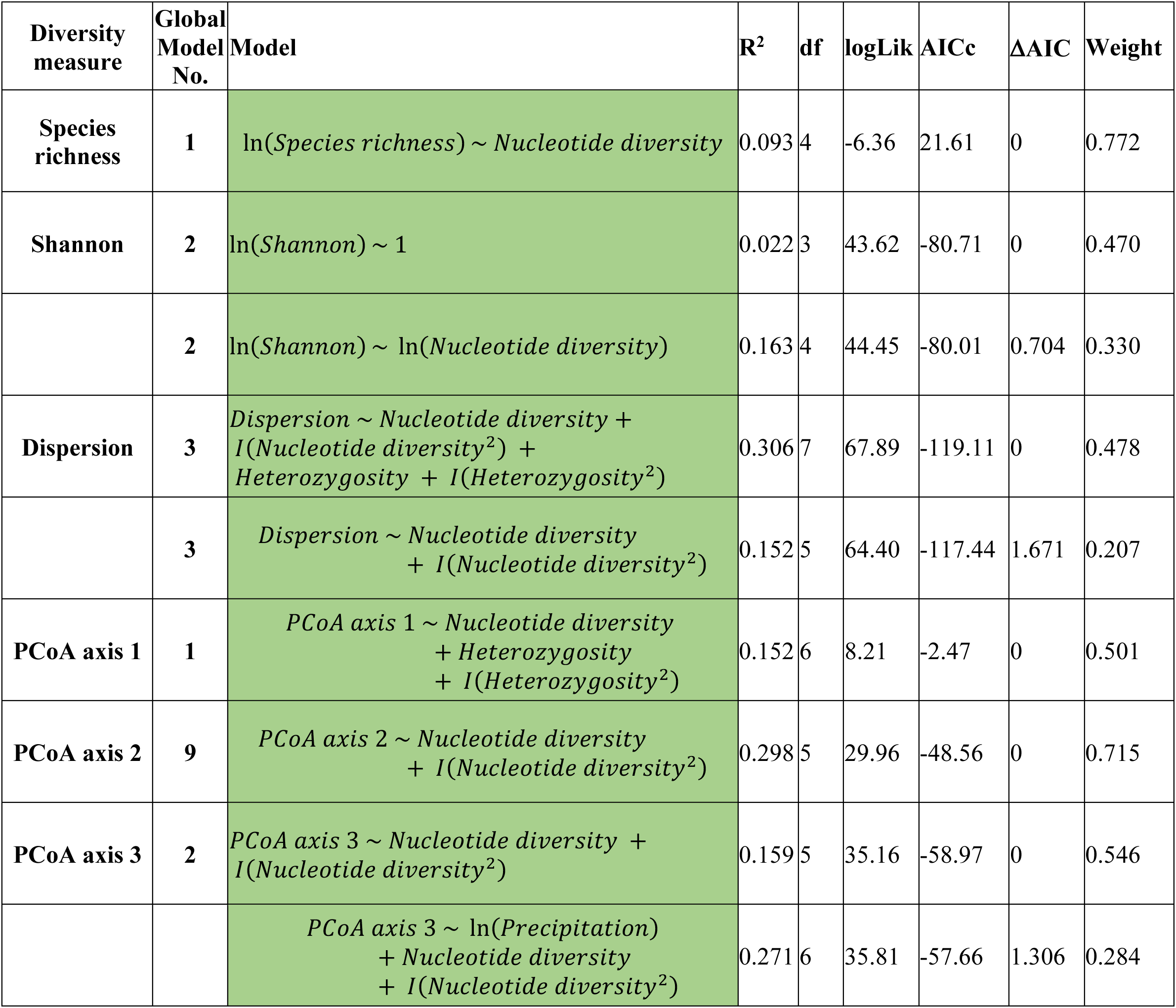
Summary of the model simplification. Models that has the lowest AIC and βAIC:< 2 can be the best simplified model.

**Table 4:**
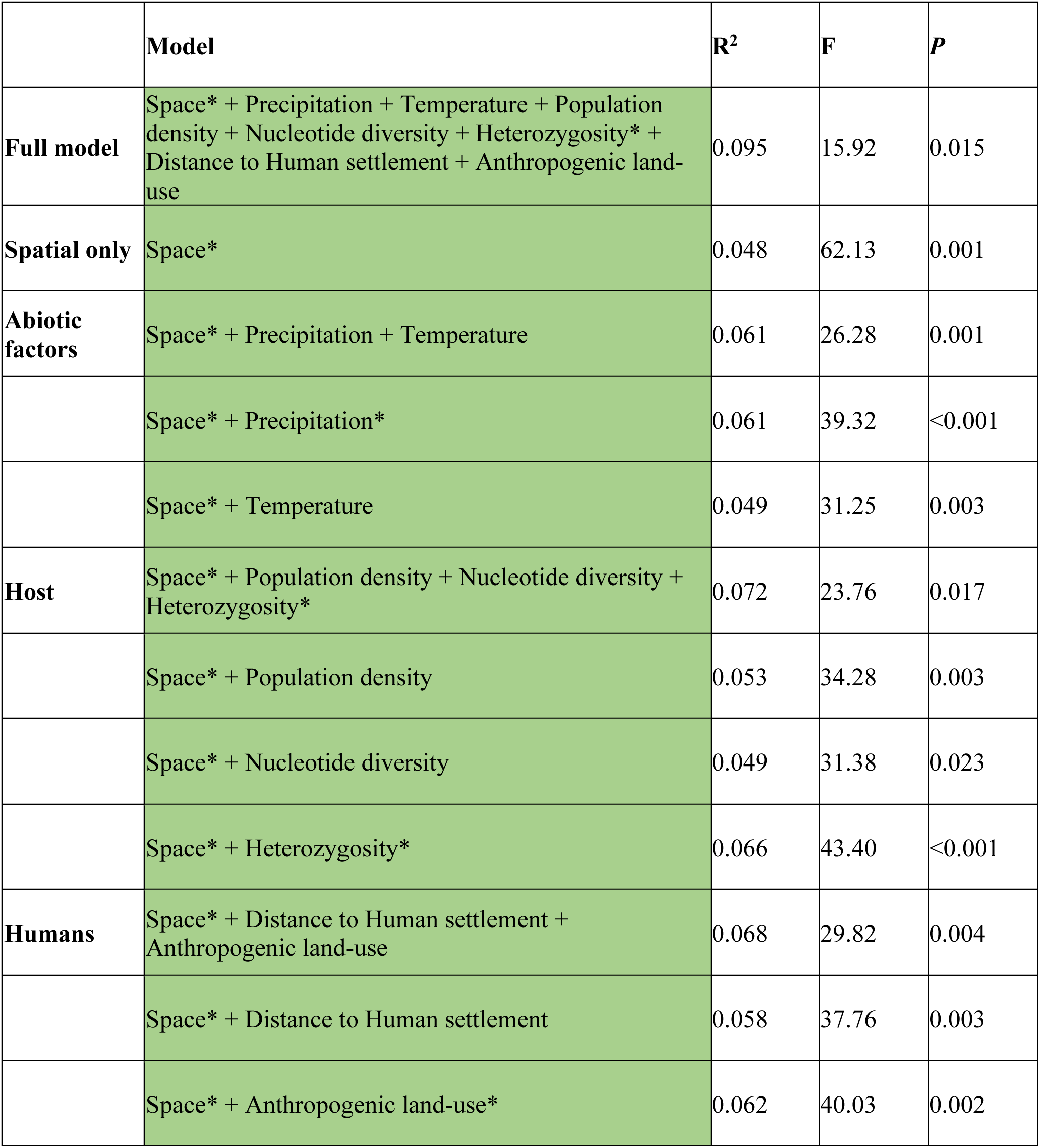
Distance based approach to understand variation in microbial diversity at landscape level (β-diversity). We summarise this using Multiple regression on distance matrices (MRM) that uses permutation tests of significance for regression coefficients and R-square. * denotes that the individual variable is significant in the model at α=0.10

|  | **Model** | **R<sup>2</sup>** | **F** | **P** |
| --- | --- | --- | --- | --- |
| **Full model** | Space* + Precipitation + Temperature + Population density + Nucleotide diversity + Heterozygosity* + Distance to Human settlement + Anthropogenic land-use | 0.095 | 15.92 | 0.015 |
| **Spatial only** | Space* | 0.048 | 62.13 | 0.001 |
| **Abiotic factors** | Space* + Precipitation + Temperature | 0.061 | 26.28 | 0.001 |
|  | Space* + Precipitation* | 0.061 | 39.32 | <0.001 |
|  | Space* + Temperature | 0.049 | 31.25 | 0.003 |
| **Host** | Space* + Population density + Nucleotide diversity + Heterozygosity* | 0.072 | 23.76 | 0.017 |
|  | Space* + Population density | 0.053 | 34.28 | 0.003 |
|  | Space* + Nucleotide diversity | 0.049 | 31.38 | 0.023 |
|  | Space* + Heterozygosity* | 0.066 | 43.40 | <0.001 |
| **Humans** | Space* + Distance to Human settlement + Anthropogenic land-use | 0.068 | 29.82 | 0.004 |
|  | Space* + Distance to Human settlement | 0.058 | 37.76 | 0.003 |
|  | Space* + Anthropogenic land-use* | 0.062 | 40.03 | 0.002 |

## Discussion

Various intrinsic host factors and extrinsic environmental factors can shape the gut microbiome residing within organisms. Studies on wild populations have yet to reach a consensus on whether host or environmental factors strongly influence the gut microbiome composition (Kartzinel et al., 2019; Knutie et al., 2019). Here we delineate the effects of host and environment on different measures of gut microbial diversity in blackbuck *A. cervicapra* by sampling individuals from a range of potential intrinsic and extrinsic factors. Contrary to our expectation, we find that not all the seven predictor variables influence blackbuck’s gut microbiome α-diversity and β-diversity. There was variation in the magnitude and direction of a predictor’s effect on microbiome diversity. Linear mixed effects modelling followed by model simplification demonstrated that measures of intrinsic host genetic diversity, rather than extrinsic environmental factors, are more important for both α- and β-diversity of blackbuck gut microbiome. The importance of host genetic diversity on the microbiome was further corroborated by multiple regression on distance matrices analyses that showed that heterozygosity, after considering background spatial structure, explains a significant variation in the gut microbial communities between individuals. These results further strengthen the concept of holobiont, where both host genetics and gut microbiota interact together to impact the evolution, ecology and health of the host organism (Simon et al., 2019).

### Blackbuck’s gut microbiome

We show that blackbuck gut is complex, with 2056 OTUs on average per individual identified across 30 known phyla. Firmicutes and Bacteroidetes were the most dominant phyla, constituting ∼75% of the total gut microbiome. They are the most common phyla in the animal gut, including humans (Bensch et al., 2023; Huttenhower et al., 2012; Ley et al., 2008; Mariat et al., 2009; Murillo, Schneider, Fichtel, et al., 2022; Xiang et al., 2020). Until recently, in humans and mice, the relative abundance of Firmicutes and Bacteroidetes are considered markers of gut dysbiosis (imbalance in the metabolic functions of gut microbiota) and obesity-related disorders (John & Mullin, 2016; Ley et al., 2006; H. Li et al., 2016; Turnbaugh et al., 2006). But the synthesis of different studies in humans examining the relationship of Firmicutes’ and Bacteroidetes’ relative abundance with dysbiosis casts doubt on the potential relationship (Knowles et al., 2019; Magne et al., 2020; Sze & Schloss, 2016; Xu et al., 2022). Nonetheless, across different animal hosts other than humans, there are small albeit significant changes in the abundance of Firmicutes and Bacteroidetes with the environment, such as food availability and captivity status (Bensch et al., 2023; Chi et al., 2019; Ley et al., 2008). These results suggest that Firmicutes and Bacteroidetes in gut microbiota across hosts in the animal kingdom are widespread and general. Cyanobacteria is the third most dominated phyla in the blackbuck gut, although it constitutes ∼5% of the microbiome. The prevalence of cyanobacteria in blackbuck’s gut could be due to the ingestion of its plant-based diet that might have facilitated the entry of cyanobacterial cells (McGorum et al., 2015; Zheng et al., 2021).

### Influence of host on the gut microbiome

We show that host genetic diversity is more important for the blackbuck gut microbiome than their environment. This result lends support to the patterns seen for larger mammalian phylogeny and their gut microbiome. Studies across different mammals suggest that the gut microbiome follows their host’s evolutionary history (Gaulke et al., 2018; Hird, 2020; Knowles et al., 2019; Ley et al., 2008; Youngblut et al., 2019). In other words, the variation in gut microbial composition is determined by the host’s phylogenetic background such that gut microbiomes converge in phylogenetically closer host species. We show that this pattern can also be recapitulated within a single mammalian species, where an increase in genetic relatedness between blackbuck individuals decreases the variation of their gut microbial composition. It is hypothesised that near universal traits among mammals, such as placenta, viviparity, mammary glands, parental care, and sociality between conspecifics, provision for intergenerational transfer of microbes through vertical and horizontal transmission (Mallott & Amato, 2021). We speculate that these drivers could also be the proximate determinants for blackbuck gut microbiome as well. Interestingly, we also find evidence of maternal inheritance of gut microbes in our results, where blackbuck mitochondrial nucleotide diversity explains considerable variation in gut microbiota. Therefore, it follows that any adverse alteration in blackbuck diversity can also have negative consequences for the gut microbiome, which in turn can compromise blackbuck fitness and population dynamics. Additionally, the population density of blackbuck and temperature could also have been important and could have contributed towards explaining the variation in the microbiome since they were highly correlated to the nucleotide diversity (Fig. S3). Further studies are required to disentangle their effects on the microbiome relative to nucleotide diversity.

### Influence of climate on the gut microbiome

We find that precipitation explains a fraction of the total variation in gut microbiome between individuals. Precipitation influences gut microbiome across various animal hosts (Orkin et al., 2018; Li et al., 2020; Baniel et al., 2021; Liu et al., 2022; Murillo, Schneider, Fichtel, et al., 2022). The putative pathways with which precipitation can influence gut microbiome are 1. Precipitation can alter the diversity of microbial communities by redistributing microbes in the environment outside the host (Bagchi et al., 2017; Maestre et al., 2015; Murillo, Schneider, Heistermann, et al., 2022); 2. Precipitation can alter dietary water intake, which can impact the host’s physiology and fitness (Heffelfinger et al., 2018; Kihwele et al., 2020); 3. Most importantly, precipitation can alter nutrient availability for the hosts, especially for herbivores in semi-arid grasslands, by regulating the plant community composition (i.e., nutrient quality) and its primary productivity (i.e., nutrient quantity) on which herbivores feed (Craine et al., 2010; Polley et al., 2013). However, further investigations and empirical data are needed to establish the pathways with which precipitation affects the gut microbiome and its host’s fitness. Unlike studies in other hosts, temperature alone did not explain any variation in the gut microbiome of blackbuck (Bestion et al., 2017; Fan et al., 2022; Kikuchi et al., 2016; J. Li et al., 2023; McMunn et al., 2022). This could be due to the narrow range of temperatures (between 25 °C and 30.5 °C) experienced by individual blackbucks at different sampling locations. While our univariate analyses did not find any effect of temperature, temperature could be an important determinant for the gut microbiome since it was highly correlated to nucleotide diversity (Fig. S3) and was not included in the multivariate analyses. Even so, our results suggest that projected variability in precipitation patterns of future climate across the range of blackbuck can significantly impact blackbuck population dynamics through gut microbiome (Kulkarni et al., 2020).

#### Influence of Humans on the gut microbiome

Human influence on animal gut microbiome is prevalent across various host species (Knutie et al., 2019; Lavrinienko et al., 2021; San Juan et al., 2020; Wasimuddin et al., 2022). Here humans can alter nutrient availability and reduce home ranges for the host through land-use change. Humans can also alter environmental microbiota, a potential source of gut microbes, either by contaminating their habitat with microplastics, xenobiotics, metals, etc., or by introducing livestock and other domestic animals into their habitat (Qin et al., 2020; Rillig et al., 2019). Although not as important as host genetic diversity, we also find that both the blackbuck population’s distance to human settlement and anthropogenic land-use of their expected home range, measures of human influence, affect α-diversity and β-diversity blackbuck gut microbiome. Distance to human settlement and anthropogenic land-use operates inversely to signify the extent of human impact on the blackbuck’s habitat. As the distance to human settlement increases, human influence decreases, and conversely, as anthropogenic land-use increases, human influence increases. This contrast was seen in their effects on gut microbiome also. As the distance to human settlement increases, we find a reduction in the microbiome α-diversity with a concomitant increase in the variation in microbial community composition between individuals or β-diversity. On the other hand, an increase in anthropogenic land-use increases α-diversity and decreases β-diversity. Overall, we find an increase in human influence homogenises (less variable) blackbuck gut microbiome. This reduction in variability can make blackbuck susceptible to diseases with negative consequences for their fitness and adaptability to any changes in their environment.

#### Research limitations and future studies

While we show host genetic diversity is strongly associated with its gut microbial diversity, there remains ∼90% unexplained variation in microbial diversity. Part of this variation could be stochastic, but we have yet to consider every determinant possible for the gut microbiome in this study. Diet is one of the essential extrinsic parameters not evaluated in this study, although we assessed its proxies, such as climate and anthropogenic influence. Given that we find a strong effect of precipitation and humans on the gut microbiome, it is plausible that diet quality and quantity can also have an influence (Frese et al., 2015; Fujimura et al., 2016; Kartzinel et al., 2019; Youngblut et al., 2019). Further, we did not account for sex differences in blackbuck. It has been shown that male blackbuck seems to move a lot between populations compared to females, resulting in biased gene flow, which can lead to alteration in the gut microbiome (Jana & Karanth, 2023; Stoffel et al., 2020). We also did not account for age and behavioural differences between male and female blackbuck, such as lekking (Bennett et al., 2016; Isvaran, 2020; Murillo, Schneider, Fichtel, et al., 2022). Another important mammalian host-related driver of gut microbiota, although neglected in host-microbe interaction studies, is the genetic variation in the host’s adaptive immune system, particularly in major histocompatibility complex (MHC), T cell and B cell (Bolnick et al., 2014; Davies et al., 2022; Fujimura et al., 2016). Compared to other host intrinsic factors immune system has more direct reciprocal interaction with host microbiota in general. For example, the impairment of pTreg cell generation in mice alters the gut microbial metagenome (Campbell et al., 2018). We acknowledge the need for more studies with diverse intrinsic and extrinsic factors to gain a more holistic understanding of the processes that regulate the gut microbiome of an animal.

#### Conservation implications for blackbuck

Blackbuck populations have heavily crashed in the past, and their number seems to be recovering now. However, the latter may not indicate their reduced susceptibility and grassland habitats’ continued conversion does not bode well for its resident species. Decisions regarding the management of blackbuck populations would nevertheless be important in the future, both for the well-being of the focal animal and other species that share the same ecosystem. Studies conducted on a single species population would lack information regarding the potential effects of environmental gradients across the species’ range. Hence, this could undermine the efficacy of conservation-based strategies for endangered or vulnerable species. Efforts like captive breeding, reintroduction, tackling obstacles like invasions by other species, and adverse effects of chemicals on native animals and plants can all immensely benefit from studies of animal microbiomes. The surge of novel interactions between life forms across the globe is associated with an increase in microbial transmission between related species and between human and non-human microbiomes (Keesing et al., 2010). Therefore, an increasing body of microbiome studies on other organisms, especially endangered species in the sub-continent, would add to our arsenal when greenlighting conservation biology and genetics practices.

## Acknowledgements and Funding

SR received a graduate fellowship from CSIR-India and was supported by a NERC Discipline Hopping (DH) for Discovery Science grant (NE/X018180/1). AJ received a graduate fellowship from MHRD-India. This work was supported by grants to PK, KI under the partnership between the Department of Biotechnology, Govt. of India, and Indian Institute of Science (DBT-IISc partnership). AJ would like to thank the state forest departments of Rajasthan, Karnataka, Andhra Pradesh, Tamil Nadu, Maharashtra and Gujarat and the village panchayat of Bhetanai in Orissa for their permission to conduct fieldwork.

## Declarations

Conflict of Interest: The authors declare no conflict of interest

Ethical approval: All sampling was non-invasive, and the animals were not stressed or harmed in any manner.

## Data Availability

The sequences for unique haplotypes from all the individuals of *A. cervicapra* assessed in this study were submitted to Genbank with Accession numbers OP794109 - OP794335.

The 16S rRNA gene sequences from the gut microbiome of all the individuals of *A. cervicapra* assessed in this study were submitted to SRA under BioProject ID: PRJNA1066065

## Author contribution

AJ and SR conceived the project; AJ conducted fieldwork and lab analyses; SR analyzed the data; SR and AJ wrote the first draft; All authors contributed to its revisions and approved the final version.

