## Supplementary information for "Gut feeling: host and habitat as drivers of the microbiome in blackbuck (*Antilope cervicapra*)"

This supplement supplementary information includes:

Additional methods

Figures. S1 to S6

Tables S1 to S3

Additional References

**Additional methods**

**Genetic analyses**

***Nucleotide diversity***

The D loop region spanned ~ 800 base pairs and might introduce errors during its full amplification and sequencing (Jana & Karanth, 2023, 2019). Further, DNA obtained from fecal samples can be highly fragmented. Therefore, to increase the chance of successful amplification, the entire sequence was divided into four overlapping regions, and primers were designed for each of the four regions. The primers were designed using the PrimerBLAST web tool, where the melting temperature (Tm) was set between 45 °C and 60 °C with a maximum Tm difference of 3 °C and primer stringency conditions were set at default. Out of all the primer pairs generated by the tool for each region, >5 pairs per region were selected for preliminary PCR analyses. The pairs were selected after accounting for the overall GC content and maximum sequence coverage of their corresponding overlapping regions of the D loop. After preliminary trials, HV1D, INT, DLF3 and DLF4 were shortlisted as primer pairs (Jana and Karanth, 2023). These primers consistently and successfully amplified their respective regions during the preliminary trials on different samples across the range. Double-stranded polymerase chain reactions (PCRs) were performed for each primer pair for all the samples with ~ 5–20 ng/ul of the extracted DNA, 1U Taq Polymerase, 2 mM of Magnesium chloride (MgCl_2_), 0.15 mM of deoxynucleotide triphosphates (dNTPs), 2ng/ml of Bovine Serum Albumin (BSA), 10 μm forward and reverse primers and milliQ water to make up a final reaction volume of 10ul. The thermocycler was set to 94 °C initial temperature (5 min), 45–50 cycles of 94 °C denaturation (30 s), 45–63 °C annealing (30 s), 72 °C extension (1 min) and 72 °C final extension (10 min). The PCR-amplified products were then run on 1% agarose gel using electrophoresis and observed under UV light to choose successfully amplified samples.

The final sequence obtained was aligned with the GenBank data to remove the risk of potential ‘numts’ (nuclear copies of mitochondrial DNA) and ensure the accuracy of the mitochondrial DNA region of blackbucks. To correct any errors in the identification of the nucleotide bases, the obtained sequences were edited and cleaned using Chromas v2.6.5 (technelysium.com.au/ chromas.html). Cleaned sequences were then aligned using the Muscle Algorithm with default parameters in Mega v7 (Tamura et al., 2011). After the final clean-up and alignment of the complete sequences, the nucleotide diversity values for each of the sampling locations were calculated using DnaSP v6.11.01 (Rozas et al., 2017). Briefly, nucleotide diversity was calculated as follows: first, for a pair of complete D loop sequences that represent two separate individuals, the frequency of nucleotide differences between the paired sequences was determined. Second, the calculated frequency was divided by the total length (in bp) of the paired sequences to estimate nucleotide differences per base position for the pair. Third, previous steps were repeated for all the possible pairs in a given sampling location. Fourth, the nucleotide diversity was calculated as the average of the nucleotide differences per base position across all the pairs of sequences in the population.

***Heterozygosity***

A microsatellite is a short segment of DNA where a few base pairs (1-6) are repeated multiple times successively at a particular location in the genome, also known as short tandem repeats. For blackbucks, there can be one or two different alleles at each microsatellite locus depending on the maternal and paternal allelic variation. Primer pairs for each of >25 locus were thoroughly tested following multiple PCR conditions in >25 randomly selected samples across all sampling locations. However, most of the primers failed to show any positive amplification during the preliminary trials. PCR was performed for each of the seven finalised primer pair for all the samples with ~ 5–20 ng/μl of the extracted template DNA, 1x Qiagen Multiplex master mix, 10 μm of fluorescence-labelled forward (FAM or 6HEX) and reverse primers, 2 ng/ml of BSA and milliQ water to make up the final 10ul volume. The thermocycler was set at 94 °C initial temperature (5 min), 50–55 cycles of 94 °C denaturation (30 s), 50–60 °C annealing (30 s), 72 °C extension (1 min) and 72 °C final extension (10 min). Extreme precautions were taken to avoid direct light exposure during the PCR reaction setup to prevent any degradation of the fluorescent-labelled primers. The PCR products were used to generate an electropherogram. Electropherogram was generated with electrophoresis of the PCR products, followed by the capture of emitted fluorescence from the fluorescence-labelled sequences and the determination of the molecular mass. For a sample, the electropherogram represents luminescence peaks that correspond to each amplified allele at each locus.

After identification of the microsatellite alleles at each of the 7 loci, quality checks were done in three steps. First, the presence of null alleles was estimated using MICROCHECKER v2.2.3 (Van Oosterhout et al., 2004). A null allele is an allele that is rendered non-functional or inactive due to a mutation. In the context of genotyping, a null allele can result in a failure to amplify or detect the allele, leading to a false negative result. Second, allelic drop-out was checked using MICROCHECKER v2.2.3. Allelic drop-out occurs when even one of the alleles fails amplification or detection during PCR, in spite of its presence in the sample. This can happen due to technical errors during the genotyping process, such as primer mismatch, poor DNA quality, variations in assay temperature, or the presence of PCR inhibitors in the sample. Our results showed that there was no evidence of null alleles and allelic drop-out in our dataset. Third, Pid/PI and Pid_sibs_/PI_sibs_ values for each population separately, both at each locus and cumulative from all loci, were calculated using GenAlEx v6.503 package (Peakall & Smouse, 2006, 2012) in Microsoft Excel 2016. This was done to accurately distinguish between individual blackbucks. Pid/PI is the probability of misidentifying two individuals as a single individual when drawn from the same randomly mating population. Pid_sibs_/PI_sibs_ is the probability of misidentifying siblings as the same individual. Pid/PI was <0.001, and Pid_sibs_/PI_sibs_ was <0.01. The values suggest that a minimum of 5 microsatellite loci were sufficient to allow for robust discrimination between two blackbucks in this study.

**Data analyses**

***Univariate analyses***

The models are linear, exponential, power, logarithm (natural log), and quadratic (Fig. 2). Each model involves different biological interpretations. 1. The linear relationship suggests that change in species diversity will be proportional to change in the magnitude of the predictor variable. 2. The exponential model indicates a decelerating relationship such that the initial increase in the magnitude of the predictor variable will be accompanied by a minimal change in the species diversity, but after a certain point, the increase in the predictor value will lead to a rapid increase or decrease in species diversity. 3. The logarithm model indicates an accelerating relationship such that a minimal increase in predictor values will lead to a rapid increase or decrease in species diversity, but after a certain point, a further increase in the predictor value will cause a minimal change in diversity. 4. The power model can fit multiple shapes facilitated by exponent ‘b’. When b is 0, the model indicates no relationship between predictor and diversity; when b is 1, the model indicates a linear relationship. If b > 1, the power model indicates an accelerating relationship and has a similar interpretation as the logarithm function. If 0 > b >1, the power model indicates a decelerating relationship and has a similar interpretation to the exponential model. 5. The quadratic model suggests species diversity is highest (or lowest due to positive quadratic term) at intermediate predictor variable values. Quadratic or unimodal fit has been documented for species diversity and productivity relationships across aquatic and terrestrial ecosystems where species diversity decreases at high productivity levels (Mittelbach et al., 2001). One of the many reasons for such a relationship could be the evolutionary trade-off between the species and its environment.

Figure S1. Sampling locations

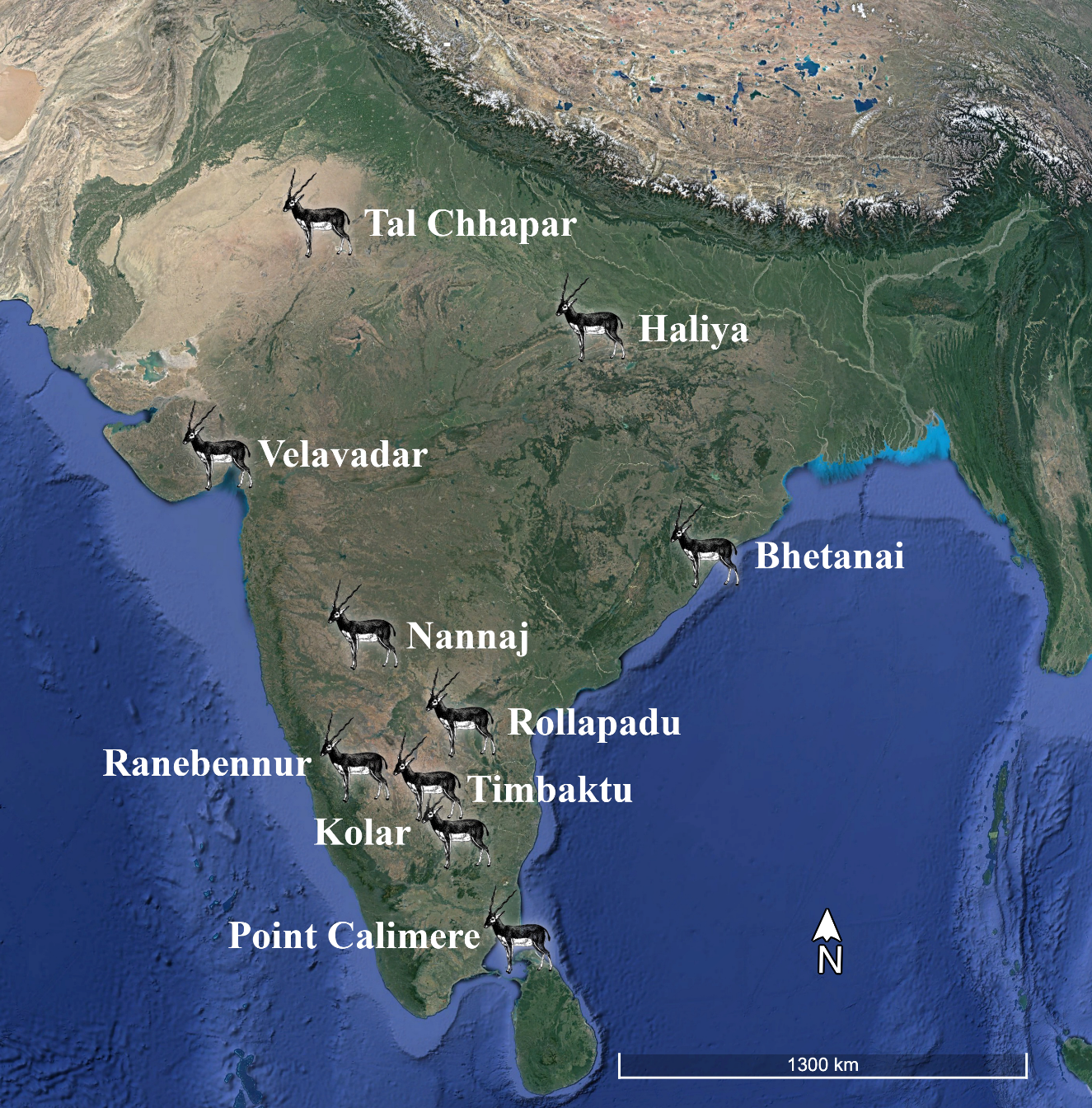

Figure S2. Summary of the predictor variables across different sampling locations and individuals

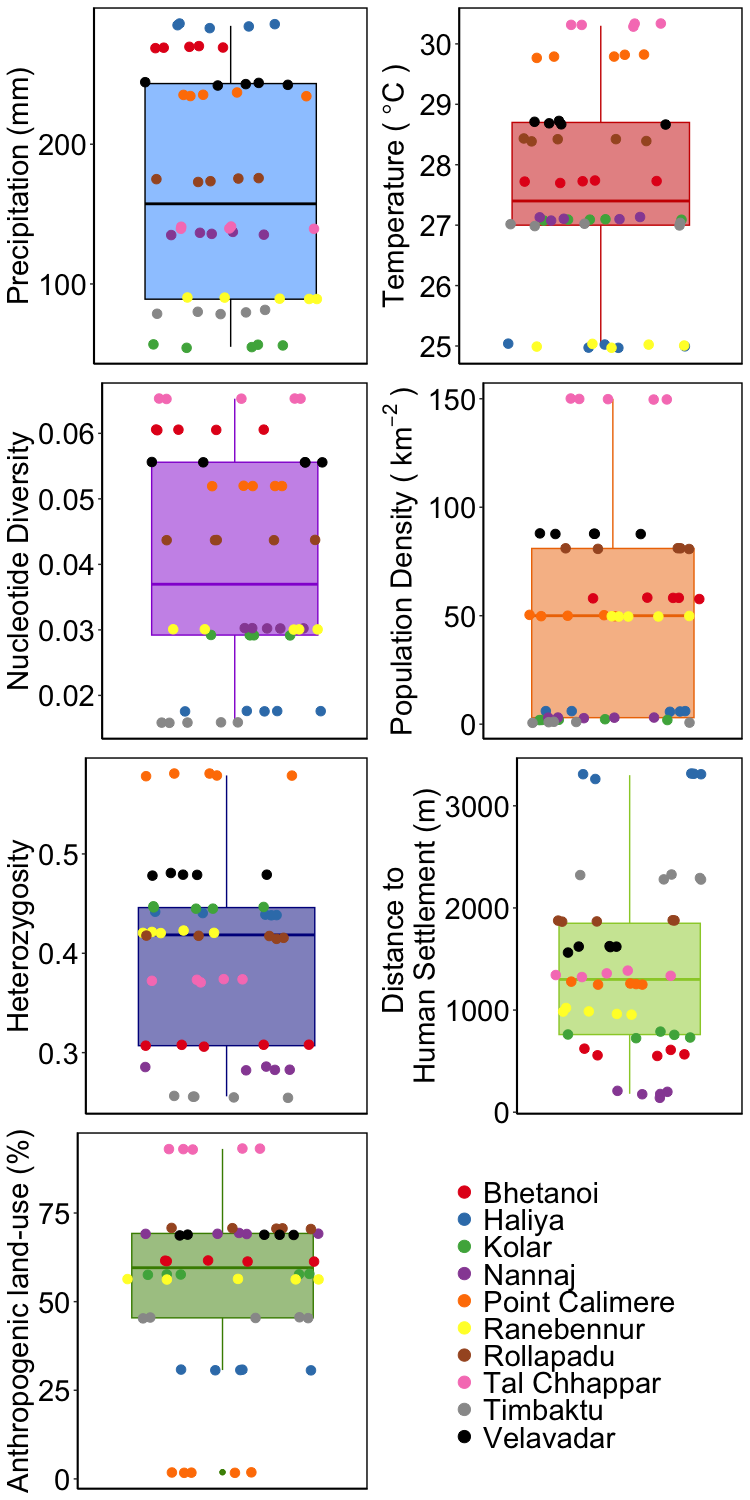

Figure S3. Correlation between explanatory variables to identify collinear variables that will be dropped for multivariate model building. We remove population density and temperature from the multivariate models.

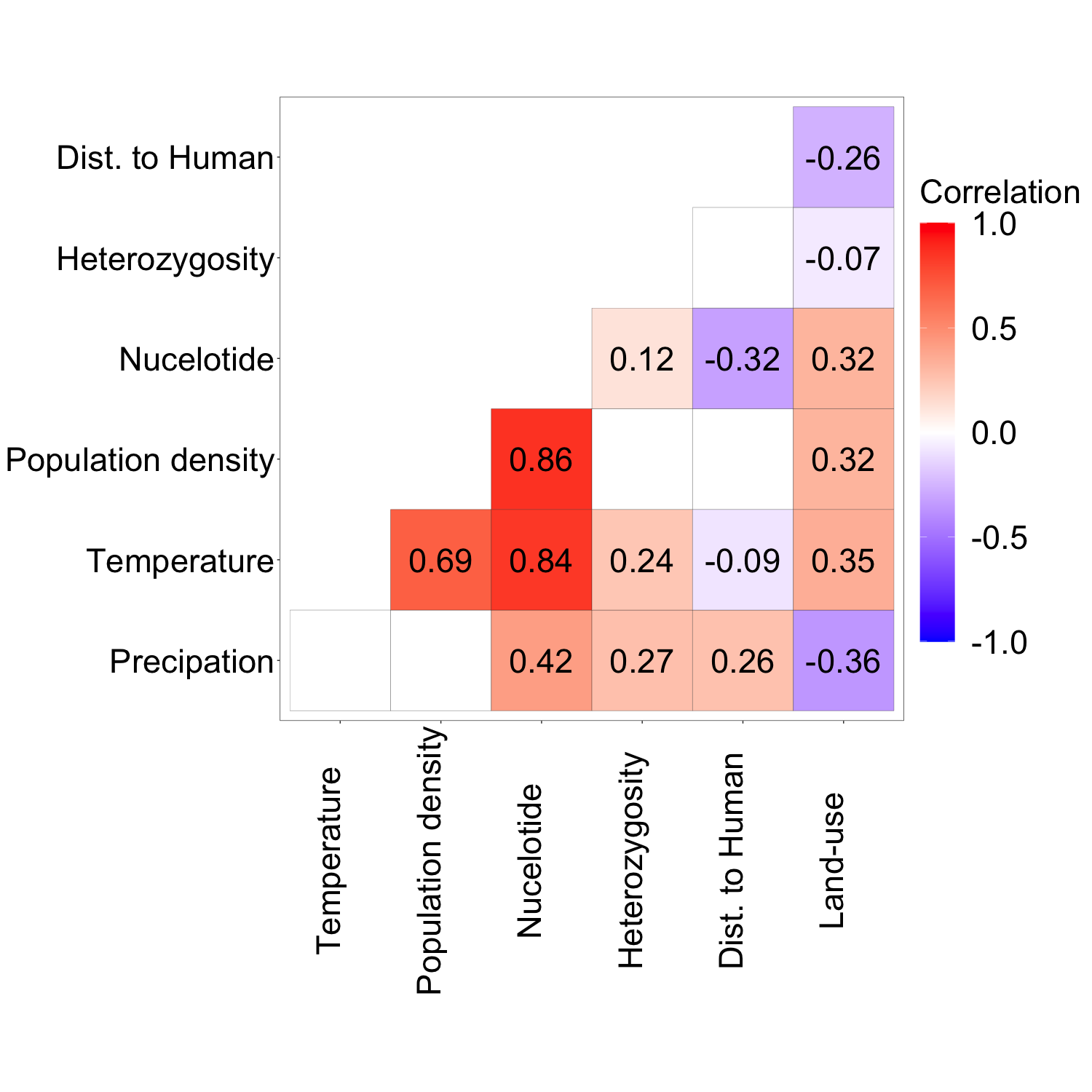

Figure S4. Relative abundance of different classes in the Firmicutes and Bacteroidetes as they are the dominant phyla across all gut microbiome Dominant phyla

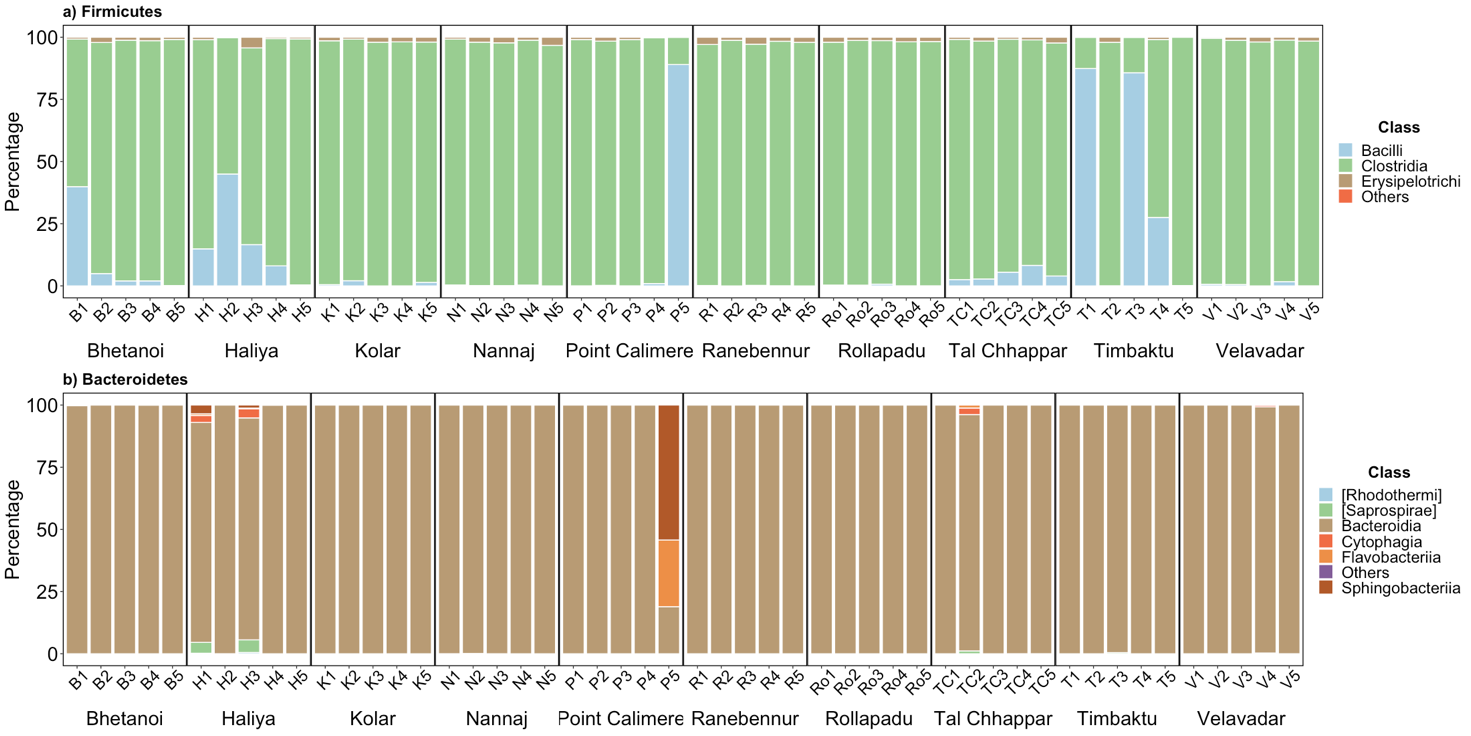

Figure S5. Visualisation of the univariate model fit for microbial diversity at the local level (α-diversity) as summarised in the Table S1.

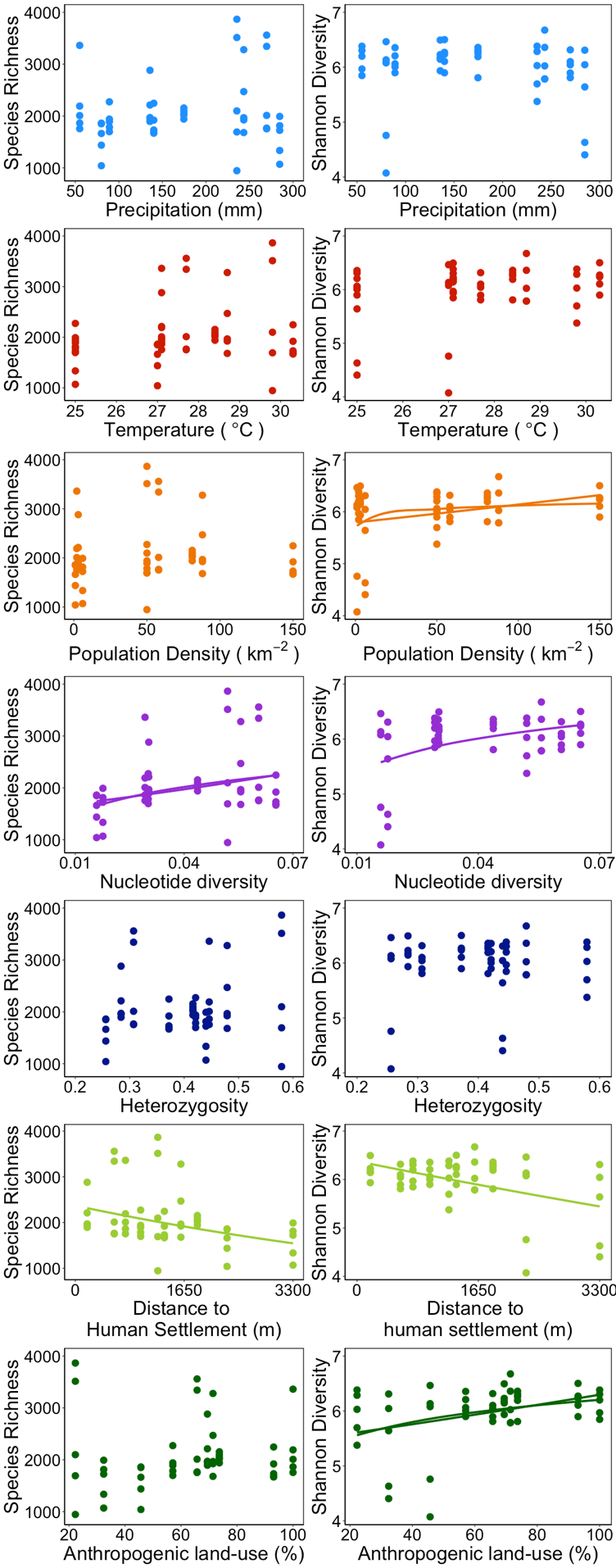

Figure S6. Visualisation of the univariate model fit for microbial diversity at landscape level (β-diversity) as summarised in Table S1.

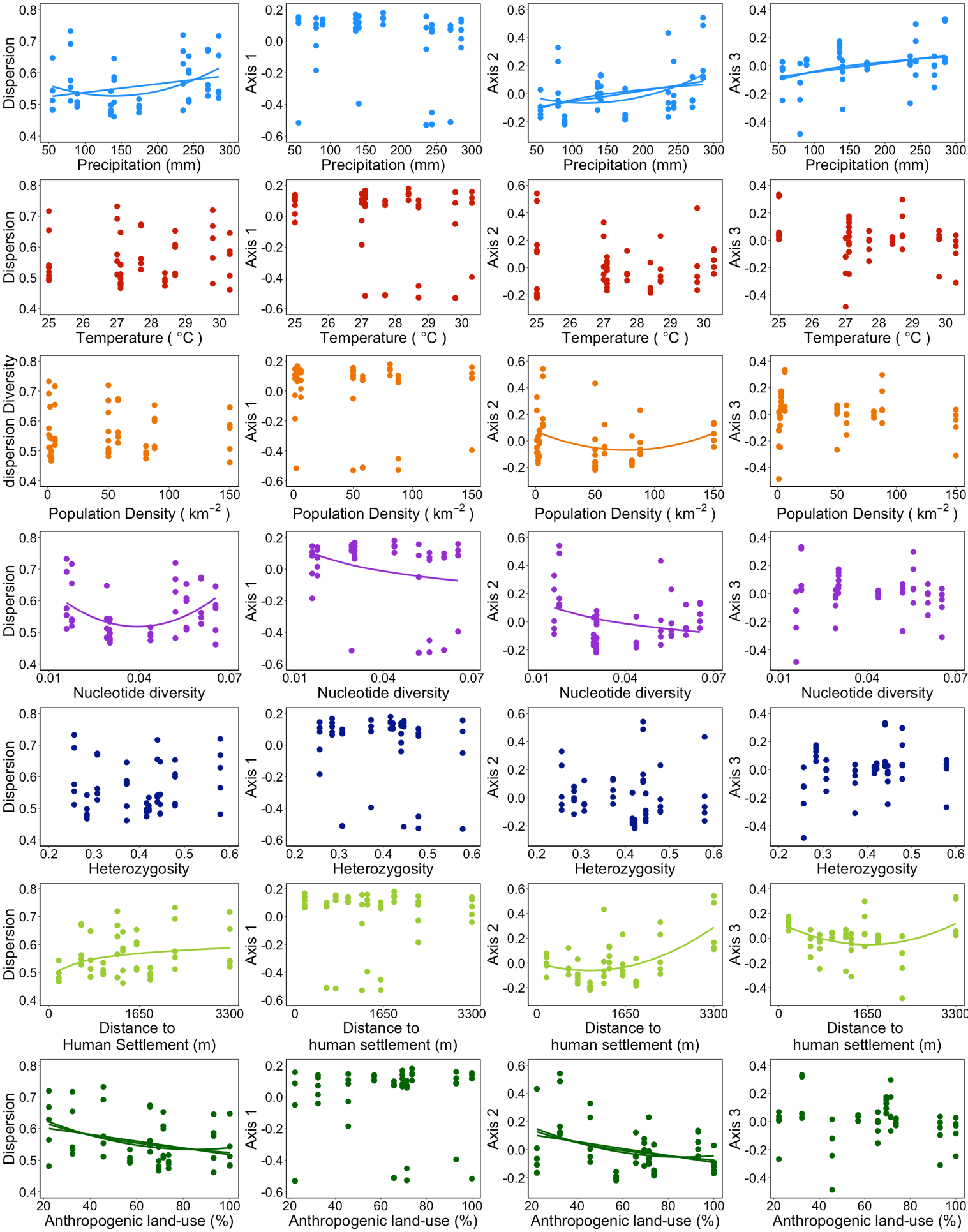

Table S1: Model fit statistics and AICc index for the different functions describing the relationship for all combinations of species diversity index and predictors. Here, species richness and Shannon represent microbial diversity at the local level (α-diversity). PCoA axis 1—3 and dispersion represent microbial diversity at the landscape level (β-diversity). The models are Linear (y=a + b×x), Exponential (y=a × e^bx^), Power (y=a × x^b^), Logarithm (y=a + b log(x)), and Quadratic (y=a + b×x + c×x^2^). AIC measures a given model's relative goodness of fit; the lower its value, the more likely it is that this model is correct. Two models with ΔAIC > 2 are substantially different. Red indicates no relationship between species diversity and predictor variables. Green indicates a significant relationship with the lowest AIC. A: accelerating relationship; D: decelerating relationship; L: Linear relationship

| **Diversity index** | **Predictor variable** | **Model** | **Parameter estimate** | **95% CI** | **R^2^** | **AIC** | **ΔAIC** | **Selected model** |
| --- | --- | --- | --- | --- | --- | --- | --- | --- |
| **Species Richness** | **Precipitation** | Null | a= 2056.68 | a= 1878.02—2235.34 |  | 645.35 | 767.12 |  |
|  |  | Linear | a=1881.40  b=1.03 | a=1455.87—2306.94  b= -1.23—3.28 | 0.017 | 646.49 | 768.26 |  |
|  |  | Exponential | a=7.54  b=0.00 | a= 7.34—7.74  b= -0.001—0.001 | 0.006 | -121.74 | 0.03 |  |
|  |  | Power | a=7.38  b=0.04 | a= 6.61—8.15  b= -0.11—0.20 | 0.006 | -121.77 | 0 |  |
|  |  | Logarithm | a=1338.97  b=143.26 | a= -336.45—3014.39  b= -189.25—475.77 | 0.015 | 646.57 | 768.34 |  |
|  |  | Quadratic | a=1652.84  b=4.39  c=-0.01 | a= 636.51—2669.18  b= -9.37—18.15  c= -0.05—0.03 | 0.022 | 648.22 | 769.99 |  |
| **Species Richness** | **Temperature** | Null | a= 2056.68 | a= 1878.02—2235.34 |  | 645.35 | 768.89 |  |
|  |  | Linear | a= -125.11  b= 79.02 | a= -3022.65—2772.42  b= -25.73—183.77 | 0.046 | 645.01 | 768.55 |  |
|  |  | Exponential | a= 6.67  b= 0.03 | a= 5.34—8.00  b= -0.02—0.08 | 0.038 | -123.41 | 0.13 |  |
|  |  | Power | a= 4.47  b= 0.94 | a= 0.09—8.85  b= -0.38—2.26 | 0.041 | -123.54 | 0 |  |
|  |  | Logarithm | a= -5326.97  b= 2226.47 | a= -14851.13—4197.19  b= -644.96—5097.90 | 0.048 | 644.88 | 768.42 |  |
|  |  | Quadratic | a= -29011.81  b= 2185.15  c= -38.25 | a= -73437.90—15414.27  b= -1048.83—5419.13  c= -96.94—20.45 | 0.079 | 645.21 | 768.75 |  |
| **Species Richness** | **Population density** | Null | a= 2056.68 | a= 1878.02—2235.34 |  | 645.35 | 768.73 |  |
|  |  | Linear | a= 2009.40  b= 0.97 | a= 1746.96—2271.84  b= -2.94—4.87 | 0.005 | 647.09 | 770.47 |  |
|  |  | Exponential | a= 7.56  b= 0.001 | a= 7.44—7.68  b= -0.001—0.002 | 0.011 | -122.02 | 1.36 |  |
|  |  | Power | a= 7.49  b= 0.03 | a= 7.33—7.65  b= -0.05—0.08 | 0.038 | -123.38 | 0 |  |
|  |  | Logarithm | a= 1870.47  b= 63.44 | a= 1522.08—2218.86  b= -38.65—165.53 | 0.032 | 645.75 | 769.13 |  |
|  |  | Quadratic | a= 1874.06  b= 10.14  c= -0.07 | a= 1578.83—2169.29  b= -0.51—20.79  c= -0.14—0.01 | 0.073 | 645.56 | 768.94 |  |
| **Species Richness** | **Nucleotide diversity** | Null | a= 2056.68 | a= 1878.02—2235.34 |  | 645.35 | 773.67 |  |
|  |  | Linear | a= 1612.44  b= 11103.50 | a= 1169.82—2055.06  b= 912.23—21294.77 | 0.091 | 642.59 | 770.9 |  |
|  |  | Exponential | a= 7.38  b= 5.15 | a= 7.18—7.58  b= 0.48—9.81 | 0.093 | -126.34 | 1.98 | **A** |
|  |  | Power | a= 8.30  b= 0.21 | a= 7.76—8.84  b= 0.05—0.38 | 0.128 | -128.32 | 0 | **A** |
|  |  | Logarithm | a= 3553.36  b= 450.40 | a= 2360.53—4746.19  b= 95.08—805.73 | 0.119 | 641 | 769.32 |  |
|  |  | Quadratic | a= 566.29  b= 74030.58  c= -780119.26 | a= -547.72—1680.30  b= 11402.42—136658.75  c= -1546822.32— -13416.21 | 0.165 | 640.32 | 768.63 |  |
| **Species Richness** | **Heterozygosity** | Null | a= 2056.68 | a= 1878.02—2235.34 |  | 645.35 | 767.43 |  |
|  |  | Linear | a= 1641.68  b= 1037.50 | a= 853.03—2430.33  b= -883.00—2958.01 | 0.024 | 646.14 | 768.21 |  |
|  |  | Exponential | a= 7.45  b= 0.34 | a= 7.09—7.82  b= -0.55—1.22 | 0.012 | -122.06 | 0.01 |  |
|  |  | Power | a= 7.71  b= 0.13 | a= 7.38—8.05  b= -0.22—0.47 | 0.012 | -122.08 | 0 |  |
|  |  | Logarithm | a= 2408.35  b= 372.37 | a= 1683.97—3132.73  b= -370.93—1115.66 | 0.021 | 646.3 | 768.38 |  |
|  |  | Quadratic | a= 2681.20  b= -4368.01  c= 6657.61 | a= -226.70—5589.10  b= -19043.89—10307.86  c= -11260.61—24575.83 | 0.035 | 647.54 | 769.62 |  |
| **Species Richness** | **Distance to Human settlement** | Null | a= 2056.68 | a= 1878.02—2235.34 |  | 645.35 | 775.08 |  |
|  |  | Linear | a= 2419.22  b= -0.26 | a= 2093.80—2744.64  b= -0.45— -0.06 | 0.125 | 640.66 | 770.39 |  |
|  |  | Exponential | a=7.78  b= -0.0001 | a= 7.62—7.92  b= -0.00022— -0.00004 | 0.153 | -129.73 | 0 | **A** |
|  |  | Power | a= 8.40  b= -0.12 | a= 7.69—9.12  b= -0.22— -0.02 | 0.1 | -126.69 | 3.03 |  |
|  |  | Logarithm | a= 3618.62  b= -222.54 | a= 2040.80—5196.44  b= -445.99—0.90 | 0.077 | 643.34 | 773.07 |  |
|  |  | Quadratic | a= 2334.30  b= -0.12  c= -0.001 | a= 1806.04—2862.56  b= -0.81—0.57  c= -0.0001—0.0001 | 0.128 | 642.48 | 772.21 |  |
| **Species Richness** | **Anthropogenic land-use** | Null | a= 2056.68 | a= 1878.02—2235.34 |  | 645.35 | 768.56 |  |
|  |  | Linear | a= 1907.68  b= 2.36 | a= 1388.93—2426.44  b= -5.35—10.08 | 0.008 | 646.96 | 770.17 |  |
|  |  | Exponential | a= 7.44  b= 0.002 | a= 7.2094—7.67842  b= -0.001—0.006 | 0.035 | -123.21 | 0 |  |
|  |  | Power | a= 7.16  b= 0.11 | a= 6.41—7.90  b= -0.08—0.29 | 0.028 | -122.85 | 0.36 |  |
|  |  | Logarithm | a= 1728.05  b= 80.98 | a= 74.78—3381.131  b= -323.99—485.96 | 0.003 | 647.18 | 770.39 |  |
|  |  | Quadratic | a= 2046.98  b= -2.99  c= 0.04 | a= 927.08—3166.88  b= -41.87—35.88  c= -0.27—0.36 | 0.01 | 648.87 | 772.08 |  |
| **Shannon** | **Precipitation** | Null | a= 5.98 | a= 5.84—6.13 |  | -64.28 | 166.61 |  |
|  |  | Linear | a= 6.12  b= -0.001 | a= 5.77—6.47  b= -0.003—0.001 | 0.015 | -63.04 | 167.85 |  |
|  |  | Exponential | a= 1.81  b= -0.001 | a= 1.74—1.87  b= -0.001—0.0002 | 0.013 | -230.89 | 0 |  |
|  |  | Power | a= 1.84  b= -0.01 | a= 1.58—2.10  b= -0.06—0.04 | 0.004 | -230.48 | 0.42 |  |
|  |  | Logarithm | a= 6.34  b= -0.07 | a= 4.95—7.74  b= -0.35—0.21 | 0.006 | -62.56 | 168.33 |  |
|  |  | Quadratic | a= 5.35  b= 0.01  c= 0.0001 | a= 4.54—6.16  b= -0.0004—0.02  c= -0.0001—0 | 0.101 | -65.6 | 165.29 |  |
| **Shannon** | **Temperature** | Null | a= 5.98 | a= 5.84—6.13 |  | -64.28 | 169.19 |  |
|  |  | Linear | a= 3.91  b= 0.08 | a= 1.52—6.29  b= -0.01—0.16 | 0.061 | -65.4 | 168.07 |  |
|  |  | Exponential | a= 1.40  b= 0.015 | a= 0.95—1.84  b= -0.002—0.03 | 0.061 | -233.42 | 0.06 |  |
|  |  | Power | a= 0.49  b= 0.39 | a= -0.97—1.95  b= -0.05—0.83 | 0.062 | -233.47 | 0 |  |
|  |  | Logarithm | a= -0.94  b= 2.09 | a= -8.77—6.90  b= -0.27—4.45 | 0.062 | -65.46 | 168.01 |  |
|  |  | Quadratic | a= -5.97  b= 0.802  c= -0.01 | a= -43.02—31.09  b= -1.90—3.49  c= -0.06—0.04 | 0.066 | -63.71 | 169.76 |  |
| **Shannon** | **Population density** | Null | a= 5.98 | a= 5.84—6.13 |  | -64.28 | 170.19 |  |
|  |  | Linear | a= 5.83  b= 0.003 | a= 5.63—6.04  b= -0.00004—0.006 | 0.076 | -66.22 | 168.25 |  |
|  |  | Exponential | a= 1.76  b= 0.001 | a= 1.72—1.79  b= 0—0.001 | 0.078 | -234.3 | 0.17 | **A** |
|  |  | Power | a= 1.74  b= 0.02 | a= 1.69—1.79  b= 0.0003—0.03 | 0.081 | -234.47 | 0 | **A** |
|  |  | Logarithm | a= 5.75  b= 0.08 | a= 5.47—6.03  b= -0.002—0.16 | 0.074 | -66.15 | 168.32 |  |
|  |  | Quadratic | a= 5.80  b= 0.001  c= -0.001 | a= 5.56—6.04  b= -0.004—0.01  c= -0.0001—0.00004 | 0.083 | -64.6 | 169.87 |  |
| **Shannon** | **Nucleotide diversity** | Null | a= 5.98 | a= 5.84—6.13 |  | -64.28 | 174.87 |  |
|  |  | Linear | a= 5.58  b= 10.08 | a= 5.22—5.95  b= 1.72—18.43 | 0.109 | -68.06 | 171.09 |  |
|  |  | Exponential | a= 1.71  b= 1.94 | a= 1.64—1.77  b= 0.38—3.49 | 0.116 | -236.42 | 2.72 |  |
|  |  | Power | a= 2.05  b= 0.08 | a= 1.88—2.23  b= 0.03—0.15 | 0.163 | -239.15 | 0 | **A** |
|  |  | Logarithm | a= 7.39  b= 0.42 | a= 6.42—8.36  b= 0.15—0.71 | 0.153 | -70.61 | 168.54 |  |
|  |  | Quadratic | a= 4.59  b= 69.58  c= -737.68 | a= 3.69—5.49  b= 18.99—120.16  c= -1356.93— -118.44 | 0.206 | -71.82 | 167.33 |  |
| **Shannon** | **Heterozygosity** | Null | a= 5.98 | a= 5.84—6.13 |  | -64.28 | 166.46 |  |
|  |  | Linear | a= 5.85  b= 0.35 | a= 5.19—6.51  b= -1.26—1.95 | 0.004 | -62.47 | 168.26 |  |
|  |  | Exponential | a= 1.75  b= 0.08 | a= 1.63—1.88  b= -0.22—0.38 | 0.006 | -230.54 | 0.19 |  |
|  |  | Power | a= 1.82  b= 0.04 | a= 1.71—1.93  b= -0.08—0.15 | 0.009 | -230.74 | 0 |  |
|  |  | Logarithm | a= 6.16  b= 0.18 | a= 5.55—6.76  b= -0.44—0.80 | 0.007 | -62.64 | 168.1 |  |
|  |  | Quadratic | a= 4.67  b= 6.48  c= -7.55 | a= 2.25—7.09  b= -5.74—18.70  c= -22.45—7.36 | 0.025 | -61.57 | 169.17 |  |
| **Shannon** | **Distance to Human settlement** | Null | a= 5.98 | a= 5.84—6.13 |  | -64.28 | 176.18 |  |
|  |  | Linear | a= 6.34  b= -0.0003 | a= 6.08—6.60  b= -0.0004— -0.0001 | 0.177 | -72.04 | 168.42 |  |
|  |  | Exponential | a= 1.85  b=-0.00001 | a= 1.80—1.90  b= -0.0001— -0.00002 | 0.185 | -240.46 | 0 | **A** |
|  |  | Power | a= 2.07  b= -0.04 | a= 1.83—2.31  b= -0.07— -0.006 | 0.106 | -235.84 | 4.62 |  |
|  |  | Logarithm | a= 7.47  b= -0.21 | a= 6.18—8.76  b= -0.39— -0.03 | 0.102 | -67.65 | 172.81 |  |
|  |  | Quadratic | a= 6.13  b= 8.86  c= 0 | a= 5.71—6.55  b= -0.0005—0.0006  c= 0—0 | 0.206 | -71.82 | 168.64 |  |
| **Shannon** | **Anthropogenic land-use** | Null | a= 5.98 | a= 5.84—6.13 |  | -64.28 | 172.96 |  |
|  |  | Linear | a= 5.48  b= 0.008 | a= 5.08—5.88  b= 0.002—0.01 | 0.132 | -69.38 | 167.85 |  |
|  |  | Exponential | a= 1.69  b= 0.001 | a= 1.62—1.77  b= 0.0004—0.003 | 0.13 | -237.24 | 0 | **A** |
|  |  | Power | a= 1.49  b= 0.07 | a= 1.25—1.73  b= 0.01—0.13 | 0.115 | -236.38 | 0.86 | **A** |
|  |  | Logarithm | a= 4.37  b= 0.40 | a= 3.08—5.66  b= 0.08—0.72 | 0.118 | -68.57 | 168.66 |  |
|  |  | Quadratic | a= 5.38  b= 0.01  c= -0.00001 | a= 4.51—6.25  b= -0.02—0.04  c= -0.0003—0.0002 | 0.134 | -67.45 | 169.78 |  |
| **Dispersion** | **Precipitation** | Null | a= 0.56 | a= 0.54—0.58 |  | -255.86 | 3.68 |  |
|  |  | Linear | a= 0.51  b= 0.0003 | a= 0.46—0.56  b= 0—0.001 | 0.079 | -257.97 | 1.57 | **A** |
|  |  | Exponential | a= -0.68  b= 0.001 | a= -0.76— -0.59  b= 0.00002—0.001 | 0.084 | -203.7 | 55.84 |  |
|  |  | Power | a= -0.88  b= 0.06 | a= -1.22— -0.53  b= -0.01—0.13 | 0.053 | -202.07 | 57.46 |  |
|  |  | Logarithm | a= 0.39  b= 0.03 | a= 0.20—0.60  b= -0.008—0.07 | 0.05 | -256.45 | 3.09 |  |
|  |  | Quadratic | a=0.61  b= -0.001  c=0.000004 | a= 0.49—0.72  b= -0.003—0.0004  c= 0—0.00001 | 0.142 | -259.54 | 0 | **D** |
| **Dispersion** | **Temperature** | Null | a= 0.56 | a= 0.54—0.58 |  | -255.86 | 0 |  |
|  |  | Linear | a= 0.42  b= 0.01 | a= 0.06—0.78  b= -0.01—0.02 | 0.012 | -254.46 | 1.40 |  |
|  |  | Exponential | a= -0.83  b= 0.01 | a= -1.45— -0.21  b= -0.01—0.031 | 0.012 | -199.94 | 55.92 |  |
|  |  | Power | a= -1.36  b= 0.23 | a= -3.40—0.68  b= -0.38—0.85 | 0.012 | -199.91 | 55.95 |  |
|  |  | Logarithm | a= 0.12  b= 0.13 | a= -1.06—1.30  b= -0.23—0.49 | 0.011 | -254.43 | 1.43 |  |
|  |  | Quadratic | a= 1.74  b= -0.09  c= 0.002 | a= -3.85—7.34  b= -0.50—0.32  c= -0.01—0.01 | 0.017 | -252.7 | 3.16 |  |
| **Dispersion** | **Population density** | Null | a= 0.56 | a= 0.54—0.58 |  | -255.86 | 0 |  |
|  |  | Linear | a= 0.56  b= 0.0001 | a= 0.53—0.59  b= -0.001—0.0004 | 0.002 | -253.94 | 1.92 |  |
|  |  | Exponential | a= -0.59  b= -0.0001 | a= -0.65— -0.53  b= -0.001—0.001 | 0.001 | -199.38 | 56.48 |  |
|  |  | Power | a= -0.59  b= -0.003 | a= -0.66— -0.51  b= -0.03—0.02 | 0.001 | -199.4 | 56.47 |  |
|  |  | Logarithm | a= 0.56  b= -0.002 | a= 0.52—0.61  b= -0.01—0.01 | 0.001 | -253.96 | 1.90 |  |
|  |  | Quadratic | a= 0.56  b= -0.0001  c=0.0000001 | a= 0.52—0.60  b= -0.001—0.001  c= -0.00001—0.00001 | 0.002 | -251.94 | 3.92 |  |
| **Dispersion** | **Nucleotide** | Null | a= 0.56 | a= 0.54—0.58 |  | -255.86 | 4.34 |  |
|  |  | Linear | a= 0.55  b= 0.25 | a= 0.50—0.60  b= -1.06—1.55 | 0.003 | -254.01 | 6.19 |  |
|  |  | Exponential | a= -0.61  b= 0.50 | a= -0.71— -0.52  b= -1.75—2.74 | 0.004 | -199.53 | 60.67 |  |
|  |  | Power | a= -0.62  b= -0.01 | a= -0.88— -0.35  b= -0.09—0.07 | 0.001 | -199.36 | 60.85 |  |
|  |  | Logarithm | a= 0.54  b= -0.01 | a= 0.39—0.69  b= -0.05—0.04 | 0.001 | -253.91 | 6.29 |  |
|  |  | Quadratic | a= 0.73  b= -10.67  c= 135.35 | a= 0.59—0.87  b= -18.36— -2.98  c= 41.23—229.48 | 0.154 | -260.21 | 0 | **D** |
| **Dispersion** | **Heterozygosity** | Null | a= 0.56 | a= 0.54—0.58 |  | -255.86 | 1.81 |  |
|  |  | Linear | a= 0.53  b= 0.07 | a= 0.43—0.63  b= -0.17—0.30 | 0.006 | -254.18 | 3.49 |  |
|  |  | Exponential | a= -0.64  b= 0.12 | a= -0.81— -0.47  b= -0.29—0.52 | 0.007 | -199.67 | 58.00 |  |
|  |  | Power | a= -0.57  b= 0.02 | a= -0.73— -0.42  b= -0.14—0.18 | 0.002 | -199.41 | 58.26 |  |
|  |  | Logarithm | a= 0.57  b= 0.01 | a= 0.48—0.66  b= -0.08—0.10 | 0.001 | -253.93 | 3.74 |  |
|  |  | Quadratic | a= 0.91  b= -1.92  c= 2.44 | a= 0.57—1.25  b= -3.63— -0.20  c= 0.34—4.54 | 0.11 | -257.67 | 0 |  |
| **Dispersion** | **Distance to Human settlement** | Null | a= 0.56 | a= 0.54—0.58 |  | -255.86 | 2.18 |  |
|  |  | Linear | a= 0.53  b= 0.00002 | a= 0.48—0.57  b= 0—0.00001 | 0.066 | -257.29 | 0.75 |  |
|  |  | Exponential | a= -0.65  b= 0.00004 | a= -0.72— -0.58  b= 0—0.0001 | 0.067 | -202.8 | 55.24 |  |
|  |  | Power | a= -0.94  b= 0.05 | a= -1.27— -0.61  b= 0.002—0.10 | 0.083 | -203.67 | 54.37 |  |
|  |  | Logarithm | a= 0.36  b= 0.03 | a= 0.17—0.55  b= 0.001—0.06 | 0.08 | -258.05 | 0 | **D** |
|  |  | Quadratic | a= 0.52  b= 0.00004  c= 0 | a= 0.45—0.58  b= -0.0001—0.00012  c= 0—0 | 0.069 | -255.42 | 2.63 |  |
| **Dispersion** | **Anthropogenic land-use** | Null | a= 0.56 | a= 0.54—0.58 |  | -255.86 | 4.61 |  |
|  |  | Linear | a= 0.62  b= -0.001 | a= 0.56—0.68  b= -0.002— -0.0002 | 0.107 | -259.54 | 0.93 | **L** |
|  |  | Exponential | a= -0.48  b= -0.002 | a= -0.58— -0.38  b= -0.003— -0.0003 | 0.105 | -204.89 | 55.58 |  |
|  |  | Power | a= -0.18  b= -0.10 | a= -0.51—0.15  b= -0.188— -0.02 | 0.122 | -205.81 | 54.66 |  |
|  |  | Logarithm | a= 0.80  b= -0.06 | a= 0.61—0.99  b= -0.11— -0.01 | 0.124 | -260.47 | 0 | **D** |
|  |  | Quadratic | a= 0.69  b= -0.004  c= 0.00002 | a= 0.57—0.82  b= -0.008—0.001  c= -0.00001—0.0001 | 0.137 | -259.26 | 1.21 | **D** |
| **PCoA axis 1** | **Precipitation** | Null | a= 0 | a= -0.06—0.06 |  | -146.57 | 1.46 |  |
|  |  | Linear | a= 0.13  b= -0.001 | a= -0.03—0.28  b= -0.002—0.0001 | 0.067 | -148.03 | 0 |  |
|  |  | Exponential | a= -1.89  b= -0.002 | a= -2.20— -1.58  b= -0.004— -0.001 | 0.164 | -67.3 | 80.73 |  |
|  |  | Power | a= -0.80  b= -0.29 | a= -2.08—0.49  b= -0.55— -0.04 | 0.129 | -65.74 | 82.29 |  |
|  |  | Logarithm | a= 0.47  b= -0.09 | a= -0.13—1.07  b= -0.21—0.03 | 0.05 | -147.11 | 0.91 |  |
|  |  | Quadratic | a= 0.03  b= 0.001  c= 0.000004 | a= -0.34—0.38  b= -0.004—0.01  c= -0.00002—0.00001 | 0.075 | -146.44 | 1.58 |  |
| **PCoA axis 1** | **Temperature** | Null | a= 0 | a= -0.06—0.06 |  | -146.57 | 1.27 |  |
|  |  | Linear | a= 0.93  b= -0.03 | a= -0.12—1.97  b= -0.07—0.004 | 0.063 | -147.81 | 0.03 |  |
|  |  | Exponential | a= -3.23  b= 0.04 | a= -5.56— -0.91  b= -0.05—0.12 | 0.02 | -61.26 | 86.58 |  |
|  |  | Power | a= -5.58  b= 1.01 | a= -13.25—2.09  b= -1.31—3.32 | 0.021 | -61.3 | 86.54 |  |
|  |  | Logarithm | a= 3.08  b= -0.93 | a= -0.36—6.51  b= -1.96—0.11 | 0.063 | -147.84 | 0 |  |
|  |  | Quadratic | a= 3.30  b= -0.21  c= 0.003 | a= -12.99—19.59  b= -1.39—0.98  c= -0.02—0.03 | 0.064 | -145.9 | 1.94 |  |
| **PCoA axis 1** | **Population density** | Null | a= 0 | a= -0.06—0.06 |  | -146.57 | 0 |  |
|  |  | Linear | a= 0.03  b= -0.001 | a= -0.07—0.12  b= -0.002—0.001 | 0.011 | -145.1 | 1.47 |  |
|  |  | Exponential | a= -2.28  b= 0.0006 | a= -2.49— -2.07  b= -0.003—0.004 | 0.004 | -60.63 | 85.94 |  |
|  |  | Power | a= -2.26  b= 0.004 | a= -2.55— -1.98  b= -0.08—0.09 | 0 | -60.5 | 86.07 |  |
|  |  | Logarithm | a= 0.06  b= -0.02 | a= -0.07—0.18  b= -0.06—0.02 | 0.022 | -145.68 | 0.89 |  |
|  |  | Quadratic | a= 0.05  b= -0.003  c= 0.00002 | a= -0.06—0.16  b= -0.006—0.002  c= -0.00001—0.00004 | 0.034 | -144.31 | 2.26 |  |
| **PCoA axis 1** | **Nucleotide diversity** | Null | a= 0 | a= -0.06—0.06 |  | -146.57 | 2.36 |  |
|  |  | Linear | a= 0.16  b= -3.87 | a= -0.01—0.32  b= -7.59— -0.15 | 0.083 | -148.93 | 0 | **L** |
|  |  | Exponential | a= -2.27  b= 0.50 | a= -2.65— -1.90  b= -8.41—9.41 | 0 | -60.51 | 88.42 |  |
|  |  | Power | a= -1.99  b= 0.08 | a= -3.07— -0.92  b= -0.24—0.40 | 0.007 | -60.75 | 88.18 |  |
|  |  | Logarithm | a= -0.41  b= -0.12 | a= -0.86—0.04  b= -0.26—0.01 | 0.067 | -148.06 | 0.87 |  |
|  |  | Quadratic | a= 0.04  b=3.17  c= -87.26 | a= -0.38—0.4  b= -20.61—26.95  c= -378.33—203.81 | 0.091 | -147.31 | 1.61 |  |
| **PCoA axis 1** | **Heterozygosity** | Null | a= 0 | a= -0.06—0.06 |  | -146.57 | 0.39 |  |
|  |  | Linear | a= 0.18  b= -0.44 | a= -0.11—0.46  b= -1.14—0.25 | 0.033 | -146.26 | 0.70 |  |
|  |  | Exponential | a= -2.23  b= -0.06 | a= -2.94— -1.53  b= -1.81—1.70 | 0 | -60.5 | 86.46 |  |
|  |  | Power | a= -2.27  b= -0.02 | a= -2.92— -1.62  b= -0.68—0.64 | 0 | -60.5 | 86.46 |  |
|  |  | Logarithm | a= -0.14  b= -0.15 | a= -0.40—0.13  b= -0.42—0.13 | 0.024 | -145.76 | 1.19 |  |
|  |  | Quadratic | a= -0.62  b= 3.69  c= -5.09 | a= -1.65—0.41  b= -1.51—8.89  c= -11.44—1.26 | 0.084 | -146.96 | 0 |  |
| **PCoA axis 1** | **Distance to Human settlement** | Null | a= 0 | a= -0.06—0.06 |  | -146.57 | 0 |  |
|  |  | Linear | a= -0.03  b= 0.00002 | a= -0.15—0.10  b= -0.0001—0.0001 | 0.005 | -144.81 | 1.76 |  |
|  |  | Exponential | a= -2.09  b= -0.0001 | a= -2.35— -1.83  b= -0.0003—0.00004 | 0.056 | -62.69 | 83.88 |  |
|  |  | Power | a= -1.80  b= -0.06 | a= -3.007— -0.60  b= -0.24—0.11 | 0.016 | -61.1 | 85.47 |  |
|  |  | Logarithm | a= 0.05  b= -0.01 | a= -0.55—0.64  b= -0.09—0.08 | 0.001 | -144.6 | 1.97 |  |
|  |  | Quadratic | a= 0.05  b= -0.0001  c= 0 | a= -0.16—0.25  b= -0.0004—0.0002  b^2^= 0—0 | 0.022 | -143.68 | 2.89 |  |
| **PCoA axis 1** | **Anthropogenic land-use** | Null | a= 0 | a= -0.06—0.06 |  | -146.57 | 0 |  |
|  |  | Linear | a= -0.06  b= 0.001 | a= -0.25—0.13  b= -0.002—0.004 | 0.009 | -145.04 | 1.53 |  |
|  |  | Exponential | a= -2.58  b= 0.01 | a= -3.02— -2.13  b= -0.002—0.01 | 0.063 | -62.95 | 83.62 |  |
|  |  | Power | a= -3.30  b= 0.26 | a= -4.77— -1.83  b= -0.101—0.61 | 0.055 | -62.66 | 83.91 |  |
|  |  | Logarithm | a= -0.27  b= 0.07 | a= -0.87—0.32  b= -0.08—0.21 | 0.018 | -145.46 | 1.11 |  |
|  |  | Quadratic | a= -0.22  b= 0.01  c= -0.0001 | a= -0.63—0.18  b= -0.01—0.02  c= -0.0002—0.0001 | 0.027 | -143.93 | 2.64 |  |
| **PCoA axis 2** | **Precipitation** | Null | a= 0 | a= -0.05—0.05 |  | -172.37 | 4.73 |  |
|  |  | Linear | a= -0.13  b= 0.001 | a= -0.25— -0.02  b= 0.0002—0.001 | 0.123 | -176.95 | 0.15 | **L** |
|  |  | Exponential | a= -3.73  b= 0.01 | a= -5.28— -2.18  b= -0.001—0.01 | 0.165 | 12.69 | 189.79 |  |
|  |  | Power | a= -7.44  b= 0.98 | a= -13.82— -1.06  b= -0.27—2.22 | 0.132 | 13.49 | 190.59 |  |
|  |  | Logarithm | a= -0.52  b= 0.10 | a= -0.97— -0.07  b= 0.01—0.19 | 0.101 | -175.71 | 1.39 | **D** |
|  |  | Quadratic | a= 0.04  b= -0.002  c= 0.00001 | a= -0.23—0.30  b= -0.01—0.002  c= 0—0.00002 | 0.16 | -177.1 | 0 | **D** |
| **PCoA axis 2** | **Temperature** | Null | a= 0 | a= -0.05—0.05 |  | -172.37 | 0 |  |
|  |  | Linear | a= 0.15  b= -0.01 | a= -0.68—0.98  b= -0.04—0.02 | 0.003 | -170.51 | 1.86 |  |
|  |  | Exponential | a= 4.54  b= -0.25 | a= -4.37—13.45  b= -0.57—0.07 | 0.132 | 13.49 | 185.86 |  |
|  |  | Power | a= 20.96  b= -7.06 | a= -8.39—50.30  b= -15.90—1.79 | 0.135 | 13.41 | 185.79 |  |
|  |  | Logarithm | a= 0.55  b= -0.17 | a= -2.18—3.29  b= -0.99—0.66 | 0.003 | -170.55 | 1.83 |  |
|  |  | Quadratic | a= 8.87  b= -0.64  c= 0.01 | a= -3.86—21.61  b= -1.57—0.29  c= -0.01—0.03 | 0.042 | -170.5 | 1.88 |  |
| **PCoA axis 2** | **Population density** | Null | a= 0 | a= -0.05—0.05 |  | -172.37 | 2.25 |  |
|  |  | Linear | a= 0.02  b= -0.001 | a= -0.05—0.10  b= -0.002—0.001 | 0.018 | -171.26 | 3.36 |  |
|  |  | Exponential | a= -2.18  b= -0.01 | a= -3.04— -1.32  b= -0.02—0.01 | 0.054 | 15.19 | 189.81 |  |
|  |  | Power | a= -2.21  b= -0.09 | a= -3.38—-1.04  b= -0.46—0.27 | 0.015 | 16.01 | 190.63 |  |
|  |  | Logarithm | a= 0.06  b= -0.02 | a= -0.04—0.16  b= -0.05—0.01 | 0.042 | -172.52 | 2.11 |  |
|  |  | Quadratic | a= 0.07  b= -0.004  c= 0.00002 | a= -0.01—0.15  b= -0.01— -0.001  c= 0—0.00004 | 0.118 | -174.63 | 0 |  |
| **PCoA axis 2** | **Nucleotide diversity** | Null | a= 0 | a= -0.05—0.05 |  | -172.37 | 12.40 |  |
|  |  | Linear | a= 0.07  b= -1.70 | a= -0.06—0.20  b= -4.66—1.26 | 0.027 | -171.75 | 13.03 |  |
|  |  | Exponential | a= -1.81  b= -16.94 | a= -3.20— -0.42  b= -49.65—15.76 | 0.062 | 15.04 | 199.80 |  |
|  |  | Power | a= -4.60  b= -0.63 | a= -8.55— -0.66  b= -1.76—0.51 | 0.07 | 14.86 | 199.63 |  |
|  |  | Logarithm | a= -0.32  b= -0.10 | a= -0.66—0.03  b= -0.20—0.01 | 0.068 | -173.9 | 10.88 |  |
|  |  | Quadratic | a= 0.61  b= -34.28  c= 403.82 | a= 0.32—0.90  b= -50.62— -17.93  c= 203.69—603.95 | 0.28 | -184.77 | 0 | **D** |
| **PCoA axis 2** | **Heterozygosity** | Null | a= 0 | a= -0.05—0.05 |  | -172.37 | 0 |  |
|  |  | Linear | a= 0.03  b= -0.08 | a= -0.19—0.26  b= -0.63—0.46 | 0.002 | -170.48 | 1.90 |  |
|  |  | Exponential | a= -3.11  b= 1.70 | a= -5.95— -0.27  b= -5.35—8.75 | 0.014 | 16.03 | 188.40 |  |
|  |  | Power | a= -1.79  b= 0.68 | a=-4.48—0.90  b= -2.02—3.37 | 0.015 | 16 | 188.38 |  |
|  |  | Logarithm | a= -0.04  b= -0.05 | a= -0.25—0.16  b= -0.26—0.16 | 0.004 | -170.58 | 1.80 |  |
|  |  | Quadratic | a= 0.42  b= -2.07  c= 2.45 | a= -0.41—1.24  b= -6.23—2.08  c= -2.62—7.52 | 0.022 | -169.47 | 2.90 |  |
| **PCoA axis 2** | **Distance to Human settlement** | Null | a= 0 | a= -0.05—0.05 |  | -172.37 | 16.67 |  |
|  |  | Linear | a= -0.14  b= 0.0001 | a= -0.22— -0.05  b= 0.0001—0.00015 | 0.23 | -183.46 | 5.58 |  |
|  |  | Exponential | a= -3.32  b= 0.001 | a= -4.59—-2.04  b= -0.0001—0.001 | 0.13 | 13.52 | 202.55 |  |
|  |  | Power | a= -5.45  b= 0.41 | a= -11.12—0.23  b= -0.36—1.19 | 0.065 | 14.97 | 204.01 |  |
|  |  | logarithm | a= -0.47  b= 0.07 | a= -0.91— -0.03  b= 0.004—0.13 | 0.087 | -174.92 | 14.12 |  |
|  |  | Quadratic | a= 0.002  b= -0.0001  c= 0 | a= -0.13—0.13  b= -0.0003—0.00004  c= 0—0 | 0.339 | -189.04 | 0 | **D** |
| **PCoA axis 2** | **Anthropogenic land-use** | Null | a= 0 | a= -0.05—0.05 |  | -172.37 | 4.26 |  |
|  |  | Linear | a= 0.16  b= -0.002 | a= 0.018—0.2  b= -0.005— -0.0004 | 0.109 | -176.12 | 0.51 | **L** |
|  |  | Exponential | a= -1.40  b= -0.02 | a= -2.94—0.14  b= -0.04—0.01 | 0.12 | 13.75 | 190.38 |  |
|  |  | Power | a= 1.00  b= -0.87 | a= -4.18—6.18  b= -2.17—0.43 | 0.099 | 14.22 | 190.85 |  |
|  |  | logarithm | a= 0.55  b= -0.14 | a= 0.11—0.98  b= -0.24— -0.03 | 0.118 | -176.63 | 0 | **D** |
|  |  | Quadratic | a= 0.31  b= -0.01  c= 0.0001 | a= 0.02—0.61  b= -0.02—0.002  c= -0.00003—0.0001 | 0.136 | -175.68 | 0.95 | **D** |
| **PCoA axis 3** | **Precipitation** | Null | a= 2.85 | a= -0.04—0.04 |  | -191.25 | 5.07 |  |
|  |  | Linear | a= -0.11  b= 0.001 | a= -0.20— -0.02  b= 0.0002—0.001 | 0.128 | -196.08 | 0.25 | **L** |
|  |  | Exponential | a= -3.86  b= 0.003 | a= -5.02— -2.69  b= -0.002—0.01 | 0.051 | 12.38 | 208.71 |  |
|  |  | Power | a= -6.27  b= 0.60 | a= -11.00— -1.55  b= -0.32—1.52 | 0.058 | 12.15 | 208.47 |  |
|  |  | Logarithm | a= -0.49  b= 0.10 | a= -0.85— -0.12  b= 0.03—0.17 | 0.132 | -196.32 | 0 | **D** |
|  |  | Quadratic | a= -0.14  b= 0.0018  c= 0.000001 | a= -0.37—0.08  b= -0.002—0.004  c= -0.00001—0.00001 | 0.13 | -194.19 | 2.13 |  |
| **PCoA axis 3** | **Temperature** | Null | a= 2.85 | a= -0.04—0.04 |  | -191.25 | 0.84 |  |
|  |  | Linear | a= 0.55  b= -0.02 | a= -0.12—1.22  b= -0.04—0.004 | 0.054 | -192 | 0.09 |  |
|  |  | Exponential | a= -1.89  b= -0.05 | a= -8.69—4.91  b= -0.30—0.20 | 0.005 | 13.83 | 205.93 |  |
|  |  | Power | a= 0.86  b= -1.23 | a= -21.51—23.22  b= -8.00—5.54 | 0.005 | 13.85 | 205.95 |  |
|  |  | Logarithm | a= 1.84  b= -0.56 | a= -0.37—4.05  b= -1.22—0.11 | 0.055 | -192.09 | 0 |  |
|  |  | Quadratic | a= 4.50  b= -0.31  c= 0.01 | a= -5.91—14.92  b= -1.07—0.45  c= -0.01—0.02 | 0.065 | -190.62 | 1.47 |  |
| **PCoA axis 3** | **Population density** | Null | a= 2.85 | a= -0.04—0.04 |  | -191.25 | 0 |  |
|  |  | Linear | a= 0.01  b= -0.0002 | a= -0.05—0.07  b= -0.001—0.001 | 0.005 | -189.5 | 1.75 |  |
|  |  | Exponential | a= -2.86  b= -0.01 | a= -3.53— -2.20  b= -0.02—0.004 | 0.059 | 12.12 | 203.37 |  |
|  |  | Power | a= -2.42  b= -0.26 | a= -3.38— -1.45  b= -0.54—0.03 | 0.104 | 10.58 | 201.84 |  |
|  |  | Logarithm | a= -0.03  b= 0.01 | a= -0.11—0.06  b= -0.02—0.03 | 0.011 | -189.78 | 1.47 |  |
|  |  | Quadratic | a= -0.01  b= 0.001  c= -0.00001 | a= -0.08—0.06  b= -0.001—0.004  c= -0.00003—0.00001 | 0.036 | -189.08 | 2.17 |  |
| **PCoA axis 3** | **Nucleotide diversity** | Null | a= 2.85 | a= -0.04—0.04 |  | -191.25 | 0 |  |
|  |  | Linear | a= 0.02  b= -0.40 | a= -0.09—0.12  b= -2.88—2.09 | 0.002 | -189.36 | 1.89 |  |
|  |  | Exponential | a= -2.71  b= -12.92 | a= -3.90— -1.52  b= -41.92—16.08 | 0.028 | 13.12 | 204.38 |  |
|  |  | Power | a= -4.86  b= -0.50 | a= -8.28— -1.45  b= -1.50—0.51 | 0.034 | 12.94 | 204.19 |  |
|  |  | Logarithm | a= 0.001  b= 0.0004 | a= -0.29—0.30  b= -0.09—0.09 | 0.00E+00 | -189.25 | 1.99 |  |
|  |  | Quadratic | a= -0.19  b= 12.26  c= -156.94 | a= -0.47—0.08  b= -3.22—27.74  c= -346.45—32.57 | 0.058 | -190.23 | 1.02 |  |
| **PCoA axis 3** | **Heterozygosity** | Null | a= 2.85 | a= -0.04—0.04 |  | -191.25 | 0.23 |  |
|  |  | Linear | a= -0.11  b= 0.28 | a= -0.30—0.07  b= -0.16—0.73 | 0.033 | -190.94 | 0.54 |  |
|  |  | Exponential | a= -2.46  b= -1.77 | a= -4.50— -0.42  b= -6.56—3.01 | 0.019 | 13.39 | 204.87 |  |
|  |  | Power | a= -3.84  b= -0.71 | a= -5.62— -2.07  b= -2.61—1.19 | 0.02 | 13.37 | 204.85 |  |
|  |  | Logarithm | a= 0.12  b= 0.13 | a= -0.05—0.29  b= -0.05—0.30 | 0.044 | -191.48 | 0 |  |
|  |  | Quadratic | a= -0.60  b= 2.80  c= -3.10 | a= -1.26—0.06  b= -0.54—6.14  c= -7.17—0.98 | 0.079 | -191.37 | 0.11 |  |
| **PCoA axis 3** | **Distance to Human settlement** | Null | a= 2.85 | a= -0.04—0.04 |  | -191.25 | 4.47 |  |
|  |  | Linear | a= -0.01  b= 0.00001 | a= -0.09—0.07  b= -0.00004—0.0001 | 0.002 | -189.36 | 6.37 |  |
|  |  | Exponential | a= -3.35  b= 0.0001 | a= -4.15— -2.56  b= -0.0004—0.001 | 0.008 | 13.76 | 209.48 |  |
|  |  | Power | a= -1.90  b= -0.19 | a= -5.33—1.53  b= -0.67—0.30 | 0.021 | 13.35 | 209.08 |  |
|  |  | Logarithm | a= 0.16  b= -0.02 | a= -0.22—0.54  b= -0.08—0.03 | 0.014 | -189.98 | 5.74 |  |
|  |  | Quadratic | a= 0.13  b= -0.0002  c= 0 | a= 0.01—0.25  b= -0.0004— -0.00005  c= 0—0 | 0.156 | -195.73 | 0 | **D** |
| **PCoA axis 3** | **Anthropogenic land-use** | Null | a= 2.85 | a= -0.04—0.04 |  | -191.25 | 0 |  |
|  |  | Linear | a= 0.05  b= -0.001 | a= -0.07—0.17  b= -0.003—0.001 | 0.014 | -189.95 | 1.30 |  |
|  |  | Exponential | a= -3.31  b= 0.002 | a= -4.59— -2.03  b= -0.02—0.02 | 0.001 | 13.96 | 205.21 |  |
|  |  | Power | a= -3.68  b= 0.12 | a= -7.67—0.31  b= -0.88—1.13 | 0.002 | 13.93 | 205.19 |  |
|  |  | Logarithm | a= 0.12  b= -0.03 | a= -0.27—0.50  b= -0.12—0.07 | 0.008 | -189.64 | 1.61 |  |
|  |  | Quadratic | a= -0.08  b= 0.004  c= -0.00004 | a= -0.34—0.18  b= -0.005—0.01  c= -0.00011—0.00003 | 0.039 | -189.24 | 2.01 |  |

Table S2: Summarising univariate model fits after MOS test for significant quadratic relationships. Red indicates no relationship between species diversity and predictor variables. Green indicates a significant relationship. Text in the green labelled box indicates the function of model fit. Functions are denoted as L: Linear, E: Exponential, P: Power, Log: Logarithm, Q: Quadratic.

|  | Species richness | Shannon | Dispersion | PCoA axis 1 | PCoA axis 2 | PCoA axis 3 |
| --- | --- | --- | --- | --- | --- | --- |
| **Precipitation** |  |  | Q |  | L, Log, Q | L, Log |
| **Temperature** |  |  |  |  |  |  |
| **Population density** |  | E, P |  |  |  |  |
| **Nucleotide diversity** | E, P | P | Q | L | Q |  |
| **Heterozygosity** |  |  |  |  |  |  |
| **Distance to Human settlement** | E | E | Log |  | Q | Q |
| **Anthropogenic land-use** |  | E, P | L, Log, Q |  | L, Log, Q |  |

Table S3: Model averaged parameter estimates and 95% confidence intervals for models with ΔAIC<2 from multivariate analyses

| **Diversity measure** | **Parameter estimate** | **95% CI (low—high)** |
| --- | --- | --- |
| **Species richness** | Intercept: 7.38  Nucleotide diversity: 5.15 | Intercept: 7.18—7.58  Nucleotide diversity: -0.21—10.50 |
| **Shannon** | Intercept: 1.89  ln(Nucleotide diversity): 0.08 | Intercept: 1.61—2.18  ln(Nucleotide diversity): 0.02—0.14 |
| **Dispersion** | Intercept: 0.95  Nucleotide diversity: -12.77  I(Nucleotide diversity^2^): 159.16  Heterozygosity: -1.61  I(Heterozyogosity^2^): 2.26 | Intercept: 0.55—1.35  Nucleotide diversity: -23.81— -1.73  I(Nucleotide diversity^2^): 25.76—292.56  Heterozygosity: -3.61—0.39  I(Heterozyogosity^2^): -0.17—4.68 |
| **PCoA axis 1** | Intercept: -0.57  Nucleotide diversity:-3.60  Heterozygosity: 4.03  I(Heterozyogosity^2^): -5.32 | Intercept: -1.58—0.44  Nucleotide diversity: -8.20—0.99  Heterozygosity: -2.14—10.20  I(Heterozyogosity^2^): -12.8—2.20 |
| **PCoA axis 2** | Intercept: 0.61  Nucleotide diversity: -34.28  I(Nucleotide diversity^2^): 403.82 | Intercept: 0.18—1.04  Nucleotide diversity: -62.41— -6.14  I(Nucleotide diversity^2^): 59.40—748.24 |
| **PCoA axis 3** | Intercept: -0.43  ln(Precipitation): 0.14  Nucleotide diversity: 12.71  I(Nucleotide diversity^2^): -171.34 | Intercept: -1.26—0.40  ln(Precipitation): 0.010 —0.28  Nucleotide diversity: -16.10—41.53  I(Nucleotide diversity^2^): -526.55—183.87 |
